# Polycomb-lamina antagonism partitions heterochromatin at the nuclear periphery

**DOI:** 10.1101/2022.04.28.489608

**Authors:** Allison P. Siegenfeld, Shelby A. Roseman, Heejin Roh, Nicholas Z. Lue, Corin C. Wagen, Eric Zhou, Sarah E. Johnstone, Martin J. Aryee, Brian B. Liau

## Abstract

The genome can be divided into two spatially segregated compartments, A and B,^1,2^ which broadly partition active and inactive chromatin states, respectively. Constitutive heterochromatin is predominantly located within the B compartment and comprises chromatin that is in close contact with the nuclear lamina.^3–5^ By contrast, facultative heterochromatin marked by H3K27me3 can span both compartments.^2–5^ How epigenetic modifications, A/B compartmentalization, and lamina association collectively maintain heterochromatin architecture and function remains unclear.^6,7^ Here we developed an approach termed Lamina-Inducible Methylation and Hi-C (LIMe-Hi-C) that jointly measures chromosome conformation, DNA methylation, and nuclear lamina positioning. Through this approach, we identified topologically distinct A/B sub-compartments characterized by high levels of H3K27me3 and differing degrees of lamina association. To study the regulation of these sub-compartments, we inhibited Polycomb repressive complex 2 (PRC2), revealing that H3K27me3 is an essential factor in sub-compartment segregation. Unexpectedly, PRC2 inhibition also elicited broad gains in lamina association and constitutive heterochromatin spreading into H3K27me3-marked B sub-compartment regions. Consistent with repositioning to the lamina, genes originally marked with H3K27me3 in the B compartment, but not in the A compartment, remained largely repressed, suggesting that constitutive heterochromatin spreading can compensate for loss of H3K27me3 at a transcriptional level. These findings demonstrate that Polycomb sub-compartments and their antagonism with nuclear lamina association are fundamental organizational features of genome structure. More broadly, by jointly measuring nuclear position and Hi-C contacts, our study demonstrates how dynamic changes in compartmentalization and nuclear lamina association represent distinct but interdependent modes of heterochromatin regulation.

## Introduction

The spatial organization of DNA within the nucleus is critically involved in fundamental cellular processes ranging from transcription to DNA repair and replication.^3,5,8^ Not surprisingly, coordinated changes in DNA conformation and positioning are a hallmark of development, and alterations to proper genome organization have been directly linked to human disease states.^9^ The development of pioneering imaging and molecular biology approaches has catalyzed a deeper understanding of genome structure and function.^1,2,10–15^ In particular, high-throughput chromosome conformation capture (Hi-C) has revealed the fundamental role that A/B compartmentalization plays in overall 3D genome architecture. A/B compartmentalization is thought to be driven by association with different nuclear landmarks as well as multivalent interactions specified by chromatin context and transcriptional activity.^16–19^ Specifically, the B compartment is highly correlated with chromatin that associates with the nuclear lamina, a meshwork of intermediate filament proteins at the inner membrane of the nuclear envelope.^2–4^ Lamina associated domains (LADs) constitute nearly 40% of the mammalian genome and are characterized by large blocks of CpG hypomethylation and enrichment in repressive histone modifications, including H3K9me2/3 and H3K27me3.^4,7,14,20–26^ H3K9me2/3 and H3K27me3 are segregated into distinct classes of LADs, consistent with their divergent roles in mediating constitutive and facultative heterochromatin, respectively.^27–29^ However, H3K27me3 is also highly enriched at the borders of compartments and LADs, raising questions about the precise role of this mark in regulating lamina association.^4,14,24,25^

Beyond its presence within LADs, H3K27me3 has important roles in the cell-type specific regulation of gene expression and genome organization across both A and B compartments.^3,27,30^ Recent studies have demonstrated that high levels of H3K27me3, often within CpG hypomethylated regions, can mediate long-range chromatin interactions spanning from DNA loops to self-associating domains.^31–38^ These H3K27me3-mediated interactions have been implicated in the regulation of key developmental genes and are distinct from canonical structures regulated by CTCF and cohesin.^31–38^ However, how Polycomb interactions and H3K27me3 impact compartmentalization, nuclear positioning, and heterochromatin organization remains to be fully resolved.^6,7^ Deconvoluting how these layers of genome organization control cell-type specific expression programs will further our understanding of Polycomb Group (PcG) proteins and the roles they play in development and disease.^39^

## Results

### LIMe-Hi-C simultaneously identifies LADs, CpG methylation, and Hi-C contacts

Motivated by the strong connection between nuclear localization and DNA topology, we developed an approach, Lamina-Inducible Methylation and Hi-C (LIMe-Hi-C), that combines detection of protein-DNA interactions at the nuclear lamina with Hi-C to superimpose spatial information on Hi-C DNA interaction data. Recent methods integrating bisulfite conversion into the Hi-C workflow have enabled the simultaneous detection of endogenous CpG DNA methylation and Hi-C contacts, suggesting that tracking lamina association through introducing exogenous cytosine methylation could be achieved.^40–42^ Specifically, we fused the GpC cytosine methyltransferase, M.CviPI, to Lamin B1, in a strategy analogous to DNA adenine methyltransferase identification (DamID) (**Fig 1a**).^14,43,44^ Detection of this GpC methylation signature can be directly integrated into a bisulfite Hi-C workflow to simultaneously map chromatin structure, lamina association, and native CpG methylation (see Methods).

**Figure 1.**
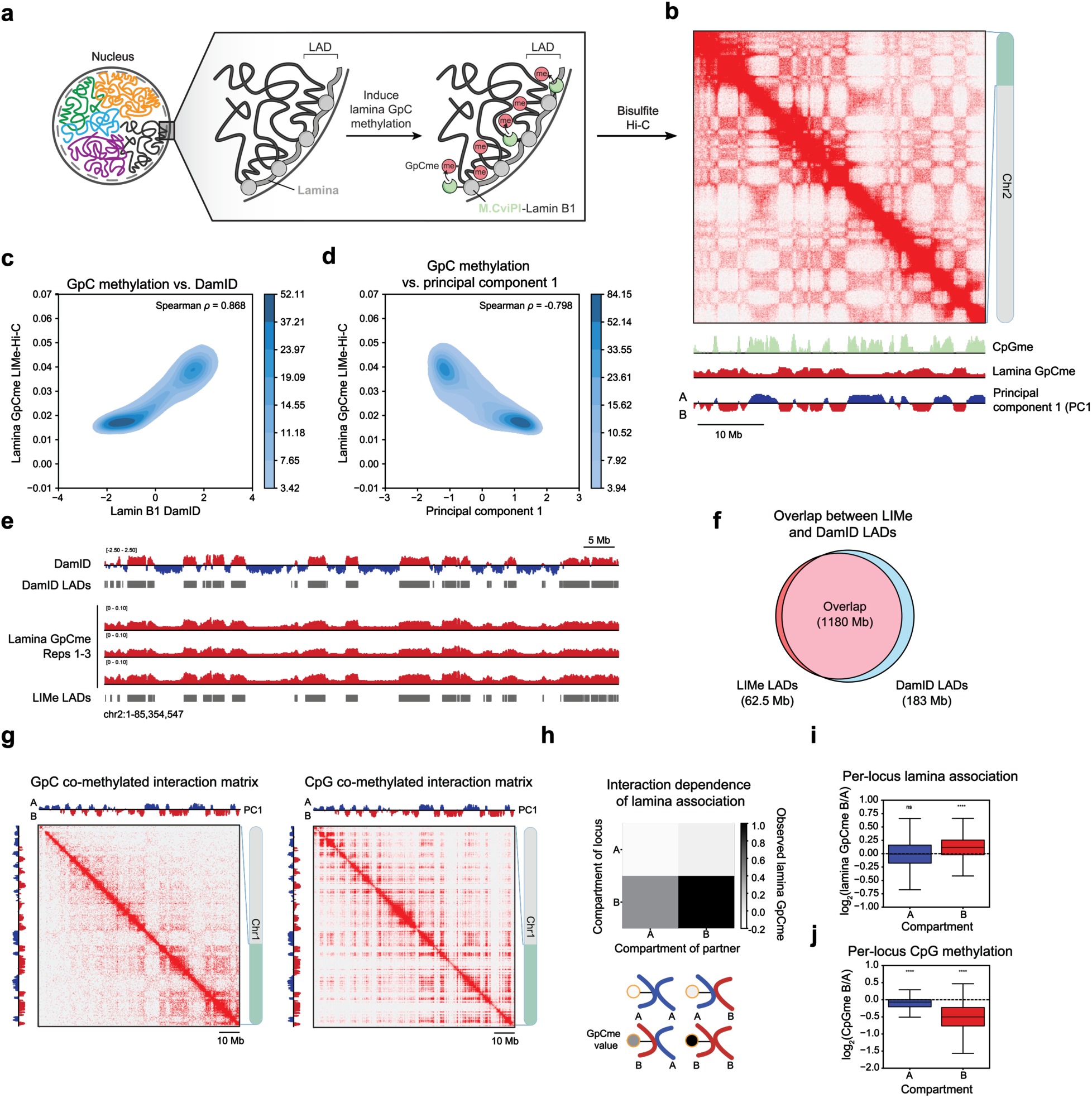
LIMe-Hi-C simultaneously identifies LADs, CpG methylation, and Hi-C contacts a) Schematic of the LIMe-Hi-C workflow. b) LIMe-Hi-C contact map for replicate 1 with lamina GpC methylation, CpG methylation, and principal component 1 at 100 kb resolution for a region on chromosome 2. c) Density heatmap comparing DamID signal (*x*-axis, Dam-Lamin B1/Dam-only enrichment) to replicate-averaged LIMe-Hi-C GpC methylation fraction (*y*-axis) across 50 kb bins. Published datasets are specified in Supplementary Table 1. d) Density heatmap comparing replicate-averaged principal component 1 (*x*-axis) to replicate-average LIMe-Hi-C GpC methylation fraction (*y*-axis) across 50 kb bins. e) Genome browser tracks depicting lamina GpC methylation signal, LIMe LADs, published DamID signal, and DamID LADs. Published datasets are specified in Supplementary Table 1. f) Venn diagram describing the genome-wide base pair overlap between LIMe LADs and published DamID LADs. Values for the set intersection and set differences are depicted. g) Example Hi-C interaction matrix for a region on chromosome 1 depicting only contacts where both reads are GpC methylated (left) or CpG methylated (right). h) Normalized average lamina GpC methylation for loci as a function of the compartment status of the loci’s interaction partner for chromosome 1 (see Methods). Curved lines represent the compartment identity of the DNA within the interaction pair. i) Boxplot for all 50 kb bins on chromosome 1 across both compartments (*x*-axis) depicting the log2 ratio of the interval’s lamina association status (*y*-axis) if it is interacting with the B versus the A compartment. Outlier points are excluded. Significance markers were calculated by a one sample one-sided t-test to determine if the mean of the population was greater than 0 and are represented as follows: ns: not significant; *: 0.01 < p ≤ 0.05; **: 0.001 < p ≤ 0.01; ***: 0.0001 < p ≤ 0.001; ****: p ≤ 0.0001. j) Boxplot for all 50 kb bins on chromosome 1 across both compartments (*x*-axis) depicting the log2 ratio of the interval’s CpG methylation status (*y*-axis) if it is interacting with the B versus the A compartment. Outlier points are excluded. Significance markers were calculated by a one sample one-sided t-test to determine if the mean of the population was less than 0 and are represented as follows: ns: not significant; *: 0.01 < p ≤ 0.05; **: 0.001 < p ≤ 0.01; ***: 0.0001 < p ≤ 0.001; ****: p ≤ 0.0001.

To perform LIMe-Hi-C, we first generated a clonal K562 cell line that expresses the Flag-M.CviPI-Lamin B1 fusion protein upon addition of doxycycline (dox). Using this cell line, LIMe-Hi-C was performed in triplicate after 24 hours of dox treatment through leveraging the bisulfite Hi-C method, Hi-Culfite,^40^ to generate Hi-C contacts with superimposed information mapping endogenous CpG methylation and exogenous GpC methylation indicative of lamina association (**Fig 1b, Fig S1a**). The bisulfite Hi-C contact data and CpG methylation profile obtained from LIMe-Hi-C are highly similar to those we obtained from performing standard *in situ* Hi-C and analyzing published whole genome bisulfite sequencing data for K562, respectively (**Fig S1b-c**).^2,40^ In addition, all three biological replicates exhibited similar profiles of GpC methylation and principal component 1 (PC1), the first eigenvector of the Hi-C Pearson correlation matrix, confirming that LIMe-Hi-C provides robust measurements of Hi-C contacts and lamina association (Spearman *ρ* > 0.95) (**Fig S1d-e**).

Consistent with the capacity of LIMe-Hi-C to profile lamina association as well as the reported connection between the B compartment and LADs, the lamina GpC methylation signal was strongly correlated with the Lamin B1 DamID signal and anticorrelated with PC1, a measure of A/B compartmentalization (**Fig 1b-d**). By contrast, in the absence of induction with dox, virtually no lamina GpC methylation signal was observed (**Fig S1f**). In addition, both the Hi-C contact heatmaps and PC1 scores are highly concordant irrespective of dox treatment, suggesting that expression of the fusion did not significantly perturb genome structure (**Fig S1f-g**). Next, we called LADs from the LIMe-Hi-C GpC methylation data after normalizing the GpC methylation signal.^45,46^ We identified 1,165 LADs with a median size of 600 kb, which significantly overlap the 1,247 LADs of median size 580 kb called by DamID (**Fig 1e-f, Fig S1h**) (**Supplementary Table 2**). Taken together, these results show that combining inducible expression of M.CviPI fused to Lamin B1 with bisulfite Hi-C enabled the robust identification of LADs, CpG methylation, and DNA-DNA contacts in a single experimental workflow.

Because LIMe-Hi-C provides single-molecule readouts of lamina association, Hi-C contact frequencies, and CpG methylation, we next investigated whether it could reveal correlations between epigenomic features that are otherwise obscured in population average measurements. Consistent with previous findings,^40^ Hi-C contact pairs on chromosome 1 are more likely than random chance to have the same CpG methylation status, suggesting a high degree of endogenous DNA methylation coordination (**Fig S1i**). To study lamina association through an analogous approach, we analyzed per-read GpC methylation data for chromosome 1 and observed a strong enrichment in contacts sharing the same lamina association status (**Fig S1i**), supporting prior models in which lamina association within single cells is correlated with bulk Hi-C interaction measurments.^4^ Taking advantage of the multi-modal data, we isolated Hi-C interaction matrices specifically containing contacts where both interaction partners possess lamina association or CpG methylation signal. The lamina-associated matrix for chromosome 1 exhibits strong enrichment of B-B interactions whereas the CpG-methylated version exhibits strong enrichment of A-A interactions (**Fig 1g**). We next explored whether a region’s interaction partner impacts its lamina association frequency and observed that B compartment regions display higher average lamina association signal when interacting with other partners from the B-versus the A compartment (**Fig 1h**). In addition, for every 50 kb region on chromosome 1, we examined the ratio of observed lamina association signals when the same region interacted with the B versus the A compartment, revealing that B compartment regions were more likely to be lamina associated when in contact with other B compartment regions (**Fig 1i**). The opposite trend is observed for CpG methylation, where B compartment loci are more likely to be CpG methylated when interacting with A compartment regions, consistent with the A compartment possessing higher levels of endogenous CpG methylation in K562 (**Fig 1j**).^34^ These data illustrate how the nuclear localization preferences and endogenous CpG methylation status of the same genomic region can vary depending on the types of loci the region is contacting. Altogether, we demonstrate that LIMe-Hi-C provides multi-modal information that can illuminate the interplay between lamina association and DNA conformation.

### Identification of topologically distinct Polycomb sub-compartments with LIMe-Hi-C

although lamina association is generally correlated with the B compartment, we observed a large degree of variability in the lamina GpC methylation signal—even for regions in the B compartment with similar PC1 values (**Fig 1d, Fig 2a, Fig S2a**). In addition, in K562 cells, the B compartment is depleted of CpG methylation while the A compartment possesses both hypo-and hypermethylated regions (**Fig S2a-b**). These observations prompted us to classify the genome into regions based on CpG methylation, lamina association, and PC1 scores. *K*-means clustering identified sub-compartment types characterized by their variance across these features (**Fig 2b, Fig S2c**) (**Supplementary Table 3**). We termed these sub-compartments Core-A, PcG-A, Core-B, and PcG-B. As described below, the Core-A and Core-B sub-compartments possess the canonical features typically associated with the A and B compartments, respectively. By contrast, the PcG-A and PcG-B sub-compartments are both enriched for the PRC2 mark, H3K27me3, and are consequently given the Polycomb group (PcG) designation.

**Figure 2.**
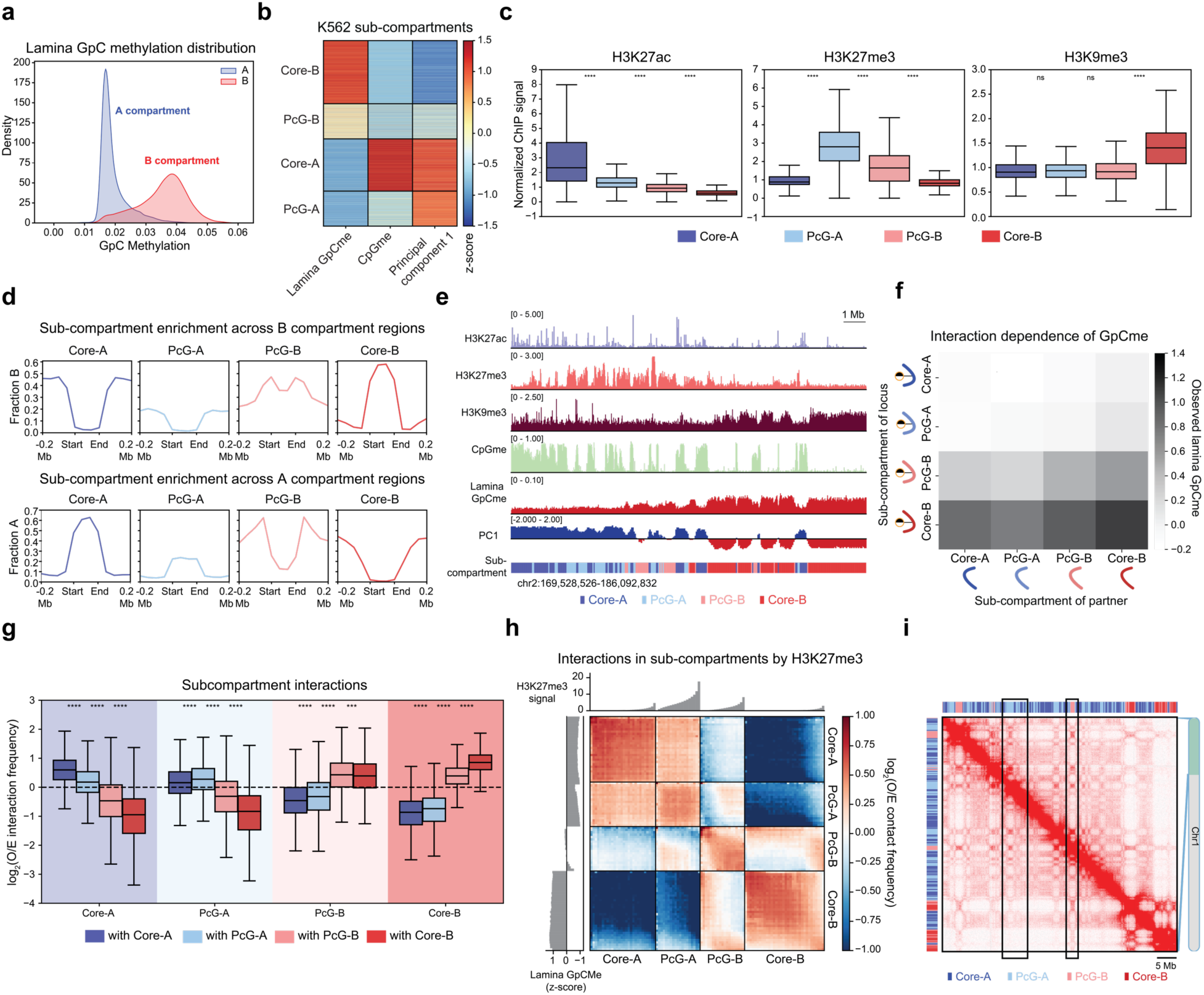
Identification of topologically distinct Polycomb sub-compartments with LIMe-Hi-C a) Density plot depicting the relative density (*y*-axis) of the average fraction lamina GpC methylation (*x*-axis) for 50 kb bins across A and B compartments in K562. The density function for each compartment is independently scaled. b) Heatmap of 50 kb genomic regions clustered into sub-compartments identified by *k*-means clustering in K562. Heatmap color represents the *z*-score transformed value for lamina GpC methylation, CpG methylation, and principal component 1 of each genomic bin. c) Boxplot showing levels of histone modifications (*y*-axis, fold-change over input/median signal) for 50 kb bins across sub-compartments (*x*-axis) in K562. Outlier points are excluded. Published datasets are specified in Supplementary Table 1. Significance markers calculated by a Mann-Whitney-Wilcoxon two-sided test are as follows: ns: not significant; *: 0.01 < p ≤ 0.05; **: 0.001 < p ≤ 0.01; ***: 0.0001 < p ≤ 0.001; ****: p ≤ 0.0001. d) Aggregate plot depicting the fraction of a specified sub-compartment type (*y*-axis) across compartment regions (*x*-axis) for the B compartment (top) and A compartment (bottom). e) Representative genome browser tracks depicting H3K27ac, H3K27me3, and H3K9me3 ChIP-seq signal along with CpG methylation (replicate 1), lamina GpC methylation (replicate 1), and principal component 1 (replicate 1) data near the *HOXD* gene cluster. The sub-compartment designation is included below. Published ChIP-seq datasets are specified in Supplementary Table 1. f) Normalized average lamina GpC methylation for loci as a function of the sub-compartment status of the loci’s interaction partner for chromosome 1 (see Methods). Curved lines represent the compartment identity of the DNA within the interaction pair. g) Boxplot showing average log2(observed/expected contact frequency) for 50 kb bin level interactions (*y*-axis) among all sub-compartments (*x*-axis). O/E denotes observed/expected. Outlier points are excluded. Significance markers calculated by a Mann-Whitney-Wilcoxon two-sided test are as follows: ns: not significant; *: 0.01 < p ≤ 0.05; **: 0.001 < p ≤ 0.01; ***: 0.0001 < p ≤ 0.001; ****: p ≤ 0.0001. h) Summary interaction heatmap ordered by sub-compartment and H3K27me3 ChIP-seq signal of the log2(observed/expected contact frequency). Replicate-averaged *z*-score lamina GpC methylation (left) and H3K27me3 levels (top) per quantile are displayed alongside the interaction heatmap. The genome was divided into 100 quantiles with the number of quantiles within a sub-compartment reflective of the relative size of the sub-compartment. O/E denotes observed/expected. i) LIMe-Hi-C contact map for K562 cells merged across the three replicates. Example PcG-A and PcG-B regions are highlighted with black boxes.

Using this classification system, A compartment regions were largely differentiated by their CpG methylation status (**Fig 2b, Fig S2c**). CpG-hypermethylated Core-A regions are highly enriched in H3K27ac while hypomethylated PcG-A regions are enriched in H3K27me3 (**Fig 2c**). By contrast, B compartment regions are chiefly differentiated by their variable association with the nuclear lamina. Core-B regions with the highest levels of lamina association are enriched in H3K9me3 (**Fig 2c**) and are located within the interior of the B compartment (**Fig 2d-e**). By contrast, PcG-B regions have PC1 scores closer to zero (less ‘B-like’), lower levels of lamina association, and enrichment in H3K27me3 (**Fig 2c, Fig S2c**). Consistent with the enrichment of H3K27me3 within PcG-A and PcG-B, these sub-compartments also displayed enrichment of proteins associated with PRC1 and PRC2 (**Fig S2d**). Upon performing H3K27me3 ChIP-seq independently, we also observed that H3K27me3 peaks overlapped to the greatest extent with the Polycomb sub-compartments (PcG-A and PcG-B) (**Fig S2e-f**). In addition, PcG-B regions are often located at compartment borders, suggesting that these regions could be involved in boundary formation (**Fig 2d**). Furthermore, B compartment regions that overlapped H3K27me3 peaks were less lamina associated than B compartment regions that did not (**Fig S2g**). Leveraging the multi-modal nature of LIMe-Hi-C, we examined the lamina association of Hi-C interaction pairs at a per-contact level. We found that the lamina association frequency of a given PcG-B locus is strongly influenced by its contact partner for regions on chromosome 1 (**Fig 2f**). In particular, B sub-compartment regions are less prone to be lamina associated when interacting with PcG-B regions compared to Core-B regions, supporting the intermediate positioning of PcG-B relative to the nuclear periphery (**Fig S2h**).

As H3K27me3 has been implicated in mediating chromatin interactions,^31–38^ we next considered if PcG-A and PcG-B regions (1) exhibit a preference for self-interactions and (2) are topologically distinct from their parental compartments. Indeed, PcG-A, PcG-B, Core-A, and Core-B regions exhibit stronger self-association within their own respective regions. Specifically, the average per-bin observed/expected Hi-C interaction frequency within a classified region is typically greater than its interaction frequency with other regions (**Fig 2g-h**). This behavior was consistent when considering replicates separately and when averaging interactions across the genome (**Fig S2i**).

Although PcG-B has a weaker propensity to self-interact, Core-B strongly self-associates (**Fig 2g-h**), suggesting that Core-B and PcG-B also segregate separately. Moreover, the interaction preferences of PcG-B regions vary depending on their baseline levels of H3K27me3. Specifically, PcG-B domains with the highest levels of H3K27me3 exhibit interaction patterns more similar to regions within PcG-A, while PcG-B domains with the lowest levels of H3K27me3 exhibit interaction patterns more similar to regions within Core-B (**Fig 2h**). Lastly, although largely partitioned into opposing compartments, PcG-A and PcG-B display moderate levels of cross-compartment Hi-C interactions, which are also observable on Hi-C contact matrices (**Fig 2i**). These data suggest that H3K27me3 marks a class of repressed chromatin embedded within the otherwise active A compartment and partitions the B compartment into two distinct sub-compartments that maintain differential contacts with the nuclear lamina and the Polycomb-repressed A compartment.

### Inhibition of EZH2 and DNMT1 differentially remodel chromatin compartmentalization

To investigate if H3K27me3 controls the structure and localization of Polycomb sub-compartments and the genome at large, we conducted LIMe-Hi-C in triplicate after chemical inhibition of EZH2, the core catalytic subunit of PRC2, with GSK343 (EZH2i).^47^ To provide a comparator to EZH2 inhibition and benchmark the robustness of the technique, we also performed LIMe-Hi-C following treatment with the non-covalent DNA methyltransferase 1 (DNMT1) inhibitor, GSK3685032 (DNMT1i).^48,49^ The inhibitors were dosed for 3 days at 1 µM and 10 µM, respectively, which led to negligible growth defects (**Fig S3a**). We verified that levels of the Flag-M.CviPI-Lamin B1 fusion were comparable across drug treatments (**Fig S3b**). Quantitative spike-in ChIP-seq demonstrated that treatment with EZH2i depleted H3K27me3 genome-wide (**Fig S3c-d**). The magnitude of a region’s change in H3K27me3 was correlated with its baseline level of H3K27me3 (**Fig S3c**), and PcG-A and PcG-B exhibited the greatest decreases in H3K27me3, consistent with these sub-compartments possessing the highest baseline levels of this modification (**Fig 2c, Fig S3e**). Treatment with DNMT1i led to an approximately 4-fold reduction in endogenous CpG DNA methylation signal as detected through LIMe-Hi-C (**Fig S3f-g**) with Core-A, the only highly methylated sub-compartment, losing the greatest amount of DNA methylation (**Fig S3h**).

At a global level, EZH2 and DNMT1 inhibition led to largely contrasting changes in compartmentalization with DNMT1 inhibition impacting compartmentalization to a larger degree than EZH2 inhibition (**Fig 3a-c, Fig S3i**). In agreement with prior studies, DNMT1 inhibition reduced genome compartmentalization on a global scale,^35,50,51^ which is reflected by the decreased ratio of intra-to inter-compartment interactions as well as broad compartment shifts, with B regions becoming more ‘A-like’ and vice versa (**Fig 3a**). At the sub-compartment level, these shifts included strengthened interactions between Core-B and both A sub-compartments (**Fig 3b, Fig 3d**). By contrast, EZH2 inhibition did not globally alter compartmentalization, but instead led to a modest strengthening of B-compartment interactions (**Fig 3a**). These effects were unexpected since Polycomb mediates chromatin compaction and has been hypothesized to be implicated in lamina association.^14,19,25,52^

**Figure 3.**
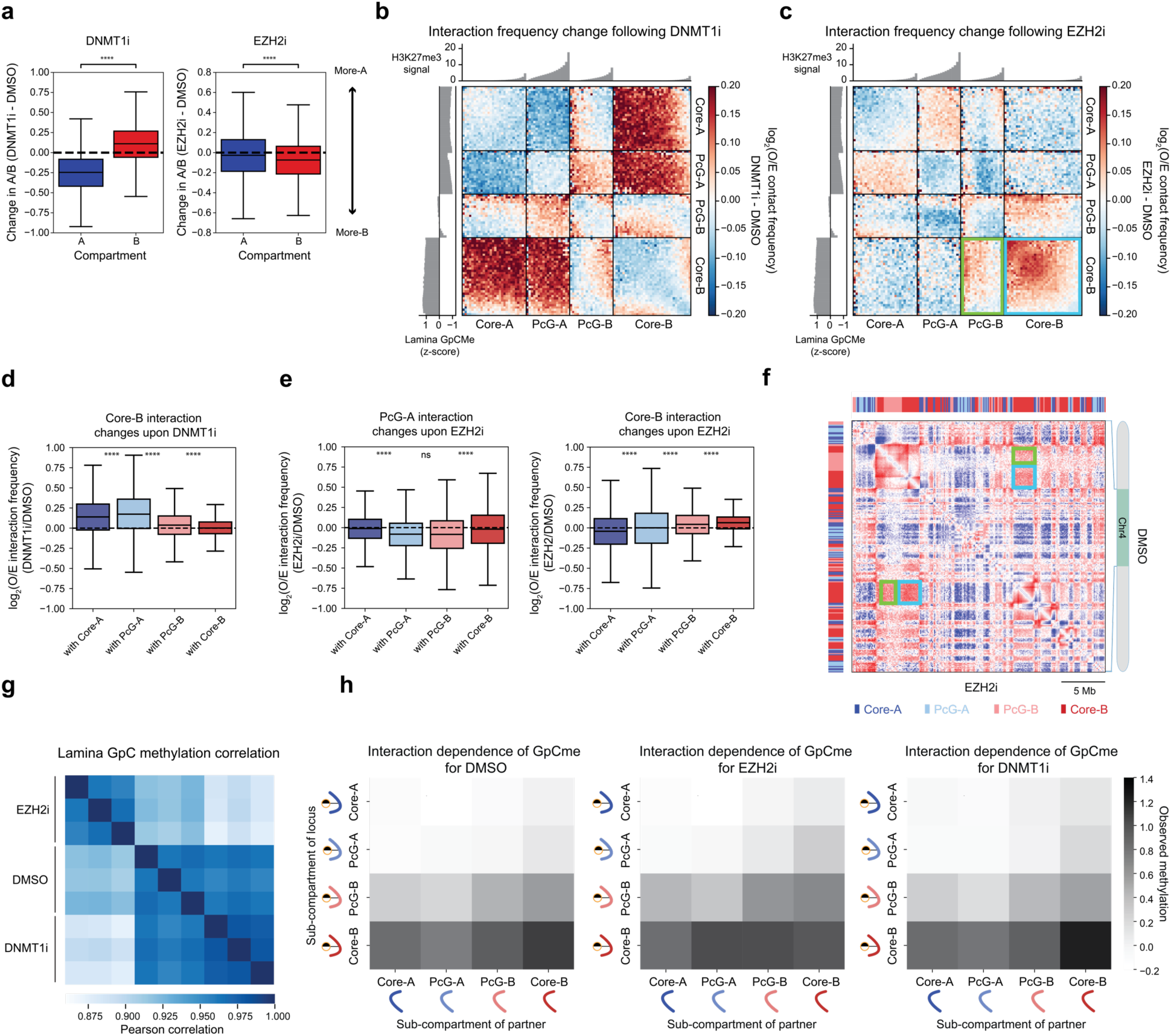
Inhibition of EZH2 and DNMT1 differentially remodel chromatin compartmentalization a) Summary boxplot plot depicting replicate average difference in log_2_(A/B) ratio for 50 kb bins (*y*-axis, log_2_(A/B ratio of observed/expected contact frequencies)) between inhibitor treatment and vehicle treatment across compartments (*x*-axis). A/B ratio represents the log of the ratio of the average observed/expected intra-chromosomal interaction frequency of a given region with A compartment bins relative to the B compartment bins (see Methods). Outlier points are excluded. Significance markers calculated by a Mann-Whitney-Wilcoxon two-sided test are as follows: ns = not significant; *0.01 < p ≤ 0.05 ** ; 0.001 < p ≤ 0.01*** ; 0.0001 < p ≤ 0.001; ****; p ≤ 0.0001. b) Summary interaction heatmap ordered by sub-compartment and baseline H3K27me3 ChIP-seq signal of the difference in the log_2_(observed/expected contact frequency) averaged across replicates between DNMT1 inhibition and vehicle treatment. Averaged baseline *z*-score lamina GpC methylation (left) and H3K27me3 (top) levels per quantile are displayed alongside the interaction heatmap. The genome was divided into 100 quantiles with the number of quantiles within a sub-compartment reflective of the relative size of the sub-compartment. O/E denotes observed/expected. c) Summary interaction heatmap ordered by sub-compartment and baseline H3K27me3 ChIP-seq signal of the difference in the log_2_(observed/expected contact frequency) averaged across replicates between EZH2 inhibition and vehicle treatment. Averaged baseline *z*-score lamina GpC methylation (left) and H3K27me3 (top) levels per quantile are displayed alongside the interaction heatmap. The genome was divided into 100 quantiles with the number of quantiles within a sub-compartment reflective of the relative size of the sub-compartment. O/E denotes observed/expected. Interactions between PcG-B and Core-B are highlighted with a green box. Interactions between Core-B and Core-B are highlighted with a blue box. d) Boxplot showing difference in log_2_(observed/expected contact frequency) of every 50 kb bin averaged across replicates (*y*-axis) for DNMT1 inhibition relative to vehicle treatment for interactions with the Core-B sub-compartment (*x*-axis). O/E denotes observed/expected. Outlier points are excluded. Significance markers calculated by a Mann-Whitney-Wilcoxon two-sided test are as follows: ns: not significant; *: 0.01 < p ≤ 0.05; **: 0.001 < p ≤ 0.01; ***: 0.0001 < p ≤ 0.001; ****: p ≤ 0.0001. e) Boxplot showing difference in log_2_(observed/expected contact frequency) of every 50 kb bin averaged across replicates (*y*-axis) for EZH2 inhibition relative to vehicle treatment for interactions with the PcG-A sub-compartment (left) and the Core-B sub-compartment (right) (*x*-axis). O/E denotes observed/expected. Outlier points are excluded. Significance markers calculated by a Mann-Whitney-Wilcoxon two-sided test are as follows: ns: not significant; *: 0.01 < p ≤ 0.05; **: 0.001 < p ≤ 0.01; ***: 0.0001 < p ≤ 0.001; ****: p ≤ 0.0001. f) Representative Hi-C observed/expected contact map comparing EZH2 inhibitor treatment to vehicle treatment. Contacts from the three replicates were pooled together. An example of increased interactions between PcG-B and Core-B is highlighted with a green box. An example of increased interactions between Core-B and Core-B is highlighted with a blue box. g) Pearson correlation heatmap of *z*-score normalized GpC methylation across 50 kb bins for LIMe-Hi-C data across replicates and inhibitor treatment conditions. h) Normalized average lamina GpC methylation for loci as a function of the sub-compartment status of the loci’s interaction partner for vehicle and inhibitor treatments for chromosome 1 (see Methods). Curved lines represent the sub-compartment identity of the DNA within the interaction pair.

To better understand the effects of EZH2 inhibition on compartmentalization, we assessed sub-compartment-specific changes in genome organization. Sub-compartment-specific analysis revealed more granular changes in interactions within the A compartment that were obscured when PcG-A and Core-A were grouped together (**Fig 3c**). Specifically, EZH2 inhibition generally decreased Hi-C self-interactions within PcG-A, which contains the highest levels of H3K27me3 (**Fig 3c, Fig 3e**). These data support the notion that H3K27me3 mediates chromatin interactions that contribute to the segregation of sub-compartments within the larger A compartment.

We next scrutinized the effects of EZH2 inhibition on the B sub-compartments. Upon EZH2 inhibition, PcG-B regions gained modest interactions with Core-B regions, which exhibited a concomitant increase in Hi-C self-interactions that drive enhanced B compartmentalization (**Fig 3e-f**). This increase in Core-B interactions suggested that H3K27me3 may be acting as a barrier to B compartment interactions. Consistent with this idea, EZH2 inhibition decreased interactions between PcG-B and PcG-A (**Fig 3c, Fig 3e**), suggesting that H3K27me3 may support interactions between these sub-compartments. Interestingly, PcG-B is heterogeneous, with regions possessing the highest levels of H3K27me3 gaining fewer B compartment interactions upon EZH2 inhibition (**Fig 3c**); it is possible that such regions may share more similarities with PcG-A. Minimal changes in CpG methylation were observed following EZH2 inhibition, suggesting that alterations in DNA methylation were not driving compartment changes (**Fig S3j-k**). Taken together, these findings suggest that H3K27me3 establishes topologically distinct sub-compartments in part by promoting interactions within PcG-A/B sub-compartments while acting as an unexpected barrier to overall B compartmentalization.

Given these observed changes in compartmentalization, we next assessed whether lamina association was substantially altered following either EZH2 or DNMT1 inhibition. Surprisingly, despite the broader compartment shifts observed upon DNMT1 inhibition (**Fig 3a, Fig S3i**), lamina association signal at a global scale was altered to a greater extent upon EZH2 inhibition (**Fig 3g**). LIMe-Hi-C per-read analysis of chromosome 1 was consistent with the more pronounced changes in lamina contacts observed upon EZH2 inhibition relative to DNMT1 inhibition. Specifically, upon EZH2 inhibitor treatment, Core-B showed higher levels of lamina association when interacting with Polycomb sub-compartments suggesting facultative heterochromatin may be repositioning from the interior to the nuclear periphery (**Fig 3h**). Altogether, these results for EZH2 and DNMT1 inhibition demonstrate that large-scale changes in lamina association are not necessarily directly coupled with changes in compartmentalization.

### PRC2 antagonizes lamina association and constitutive heterochromatin spreading

Prompted by the unanticipated observation that EZH2 inhibition strengthened B compartment interactions and altered lamina-associated contacts, we explored sub-compartment-specific changes in lamina association. Both Polycomb sub-compartments preferentially gained lamina association upon EZH2 inhibition (**Fig 4a**). Regions within PcG-B that lost the most H3K27me3— as determined by quantitative ChIP-seq—became the most lamina-attached (**Fig 4b**). Notably, this trend is not observed for PcG-A (**Fig 4b**). Furthermore, PcG-B regions that gained the most lamina association also became more B-like, although the trend is not observed for regions gaining intermediate lamina attachment (**Fig 4c**). This observation is consistent with our findings that changes in compartmentalization are related to but not always directly linked to changes in lamina association when comparing the effects of the DNMT1 and EZH2 inhibitor treatments. This shifting of PcG-B to the periphery is also evidenced by examining the change in lamina GpC methylation rank between EZH2 inhibitor and vehicle treatment (**Fig 4d)**. By contrast, no increase in lamina attachment for PcG-B regions was observed upon DNMT1 inhibition (**Fig S4a**). Thus, despite prior models indicating that in certain contexts Polycomb may mediate lamina association and cooperate with constitutive heterochromatic factors such as HP1,^3,25,53–55^ our findings suggest that H3K27me3 antagonizes direct lamina contact and interferes with B compartment interactions adding another layer of complexity to these models. Such antagonism is exemplified near the key developmental *HOXA* and *HOXD* loci, where neighboring PcG-B domains enriched in H3K27me3 gain lamina association and in some cases modest B compartment interactions following EZH2 inhibition (**Fig 4e, Fig S4b**).

**Figure 4.**
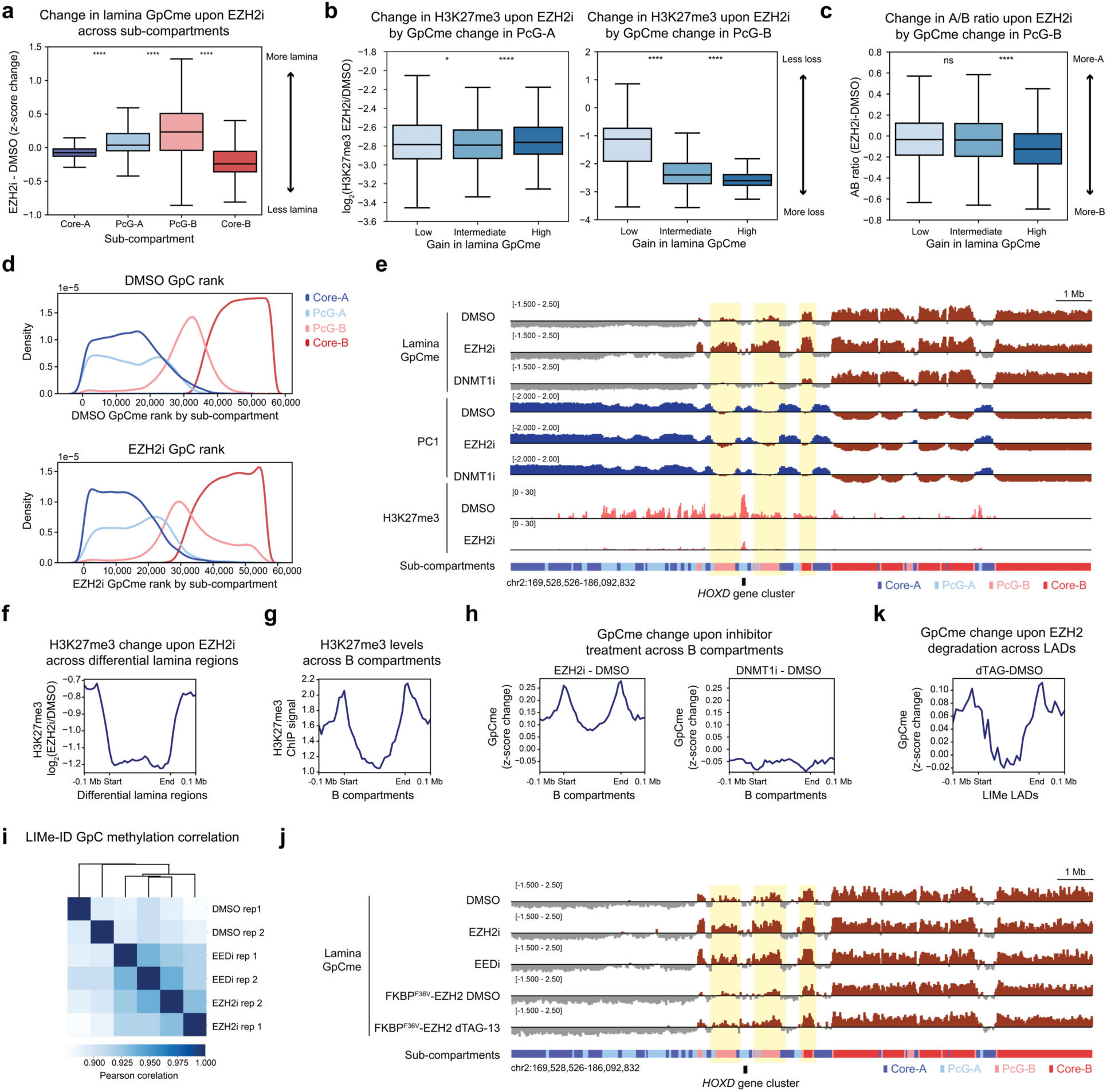
PRC2 antagonizes lamina association a) Boxplot showing the difference in *z*-score normalized GpC methylation between EZH2i and vehicle treatment for 50 kb bins (*y*-axis) across sub-compartments (*x*-axis). Outlier points are excluded. Significance markers calculated by a Mann-Whitney-Wilcoxon two-sided test are as follows: ns: not significant; *: 0.01 < p ≤ 0.05; **: 0.001 < p ≤ 0.01; ***: 0.0001 < p ≤ 0.001; ****: p ≤ 0.0001. b) Boxplot showing log_2_ fold-change in H3K27me3 levels between EZH2i and vehicle treatment for 50 kb bins (*y*-axis) segregated into three equally sized quantiles by change in *z*-score normalized lamina GpC methylation (*x*-axis) for PcG-A domains (left) and PcG-B domains (right). Outlier points are excluded. Significance markers calculated by a Mann-Whitney-Wilcoxon two-sided test are as follows: ns: not significant; *: 0.01 < p ≤ 0.05; **: 0.001 < p ≤ 0.01; ***: 0.0001 < p ≤ 0.001; ****: p ≤ 0.0001. c) Boxplot showing change in log_2_(A/B) ratio between EZH2i and vehicle treatment for 50 kb bins (*y*-axis) segregated into three equally sized quantiles by change in *z*-score normalized lamina GpC methylation (*x*-axis) for PcG-B domains. Outlier points are excluded. Significance markers calculated by a Mann-Whitney-Wilcoxon two-sided test are as follows: ns: not significant; *: 0.01 < p ≤ 0.05; **: 0.001 < p ≤ 0.01; ***: 0.0001 < p ≤ 0.001; ****: p ≤ 0.0001. d) Density plot depicting density (*y*-axis) of average lamina GpC methylation bin rank value across replicates (*x*-axis) for DMSO treatment (top) and EZH2 inhibition (bottom) for 50 kb genomic bins colored by sub-compartment designation. e) Genome browser tracks of *z*-score normalized lamina GpC methylation levels, principal component 1, and H3K27me3 ChIP-seq signal averaged across the replicates for LIMe-Hi-C and quantitative ChIP-seq data near the *HOXD* locus. f) Aggregate profile plot of log_2_ fold-change H3K27me3 ChIP-seq signal between EZH2 inhibition and vehicle treatment (*y*-axis) across regions gaining lamina contact upon EZH2 inhibition (*x*-axis) (see Methods). g) Aggregate profile plot of baseline H3K27me3 ChIP-seq signal for vehicle treatment (*y*-axis) across B compartment regions (*x*-axis). h) Aggregate profile plots depicting change in *z*-score normalized lamina GpC methylation (*y*-axis) across B compartment regions (*x*-axis) for inhibitor treatments relative to vehicle treatment. i) Clustered Pearson correlation heatmap of *z*-score normalized GpC methylation across 250 kb bins for LIMe-ID data across replicates and inhibitor treatment conditions. j) Genome browser tracks of *z*-score normalized lamina GpC methylation levels averaged across replicates for LIMe-ID inhibitor treatments and EZH2 degradation near the *HOXD* locus. k) Aggregate profile plots depicting change in *z*-score normalized lamina GpC methylation (*y*-axis) across LIMe LADs (*x*-axis) for EZH2 degradation relative to vehicle treatment for the LIMe-ID data.

To explore PRC2-lamina antagonism further, we directly called differential lamina-associated regions between EZH2i and vehicle treatment, which led to the identification of 912 regions of at least 150 kb in size that gained lamina contact (average size 301.7 kb) (**Supplementary Table 4**). Utilizing the same statistical thresholds, only 236 regions (average size 176.3 kb) were identified that lost contact with the lamina. Consistent with our prior analysis (**Fig 4b-d**), the regions that gained lamina-association significantly overlapped PcG-B and were highly depleted of H3K27me3 by EZH2 inhibition (**Fig 4f, Fig S4c-d**). By contrast, regions that lost lamina association were observed to predominantly overlap Core-B (**Fig S4e**) but comprised a significantly smaller proportion of the genome compared to regions that gain lamina association and may have been identified due to the z-score standardization employed. Given that H3K27me3 is enriched at LAD and compartment borders (**Fig 4g, S4f-g**),^14,25^ we considered whether the gains in lamina association preferentially occur at these borders versus the interior of the domains. We observed that regions gaining lamina association following EZH2 inhibition, but not DNMT1 inhibition, reside within closer proximity to both LAD and compartment borders than would be expected by random chance (**Fig S4h-i**). Moreover, upon EZH2 inhibition, lamina association substantially increased at the borders of both LADs and compartments (**Fig 4h, Fig S4j**). This trend was also evident when comparing changes in lamina association within H3K27me3 peaks at LAD borders versus within those not near LAD borders in the A compartment (iLADs) (**Fig S4k**).

To further evaluate PRC2-lamina antagonism, we assessed the effects of different PRC2 perturbations on lamina association. Instead of using the full LIMe-Hi-C workflow, we directly conducted lamina profiling by isolating and fragmenting genomic DNA before performing bisulfite conversion followed by library preparation. This approach, which we term LIMe-ID, reveals LADs and CpG methylation with less sequencing depth required and could prove to be highly relevant to study their interdependence, especially given the observation that DNA hypomethylated domains in some cancers overlap with LADs (**Fig S5a-b**).^22,24^ We first performed LIMe-ID in a different K562 clonal line that expresses the M.CviPI-LaminB1 fusion after a 72 h treatment with vehicle, GSK343 (1 µM), or EED226 (5 µM), an orthogonal PRC2 inhibitor that targets EED and blocks allosteric activation of EZH2.^56,57^ At a global level EED and EZH2 inhibitor treatments led to lamina GpC methylation signatures more similar to one another than to vehicle treatment (**Fig 4i**). In line with our previous results, lamina association increased within PcG-B upon EED and EZH2 inhibition (**Fig S5c-e**). Consistent with these changes, an increase in lamina association both at LAD borders and within regions previously determined by LIMe-Hi-C to gain lamina contact upon EZH2 inhibition was observed (**Fig S5f-g**). As before, there were minimal changes in CpG methylation for both inhibitor treatments (**Fig S5h**).

To test the effects of acute EZH2 depletion on lamina positioning, we next generated a K562 clonal cell line with an inducible degradation tag (dTAG)^58^ fused to the endogenous N-terminus of EZH2 using CRISPR/Cas9 (**Fig S5i**). Addition of the dTAG-13 ligand led to depletion of EZH2 and loss of H3K27me3 after 72 h (**Fig S5i**). LIMe-ID after acute depletion of EZH2 revealed gains in lamina association for Polycomb sub-compartments at levels similar to chemical inhibition of PRC2 (**Fig 4j, Fig S5j-l**). Furthermore, these changes were enriched at LAD borders and correlated with H3K27me3 loss (**Fig 4k, Fig S5m-n)** suggesting that H3K27me3 depletion is responsible for lamina repositioning. Collectively, these data support the notion that H3K27me3 antagonizes lamina association, particularly at the boundaries of LADs and compartments that are enriched in this modification.

We posited that H3K27me3-lamina antagonism may insulate subsets of facultative heterochromatin from the constitutive heterochromatin of the nuclear lamina environment. To assess this idea, we performed quantitative spike-in ChIP-seq for H3K9me3. Paralleling changes in lamina association, PcG-B gained the most H3K9me3 following EZH2 inhibition. In addition, increases in H3K9me3 were correlated with increases in lamina association within PcG-B (**Fig 5a-b**). Furthermore, PcG-B regions that gained the most H3K9me3 lost the most H3K27me3 (**Fig 5c**). Spreading of H3K9me3 into former H3K27me3 domains was also observed at the borders of B compartments and LADs as well as at Polycomb domains near the *HOXD* and *HOXA* loci (**Fig 5d-e, Fig S6a-b**). Altogether, these results support a model where H3K27me3 at compartment borders prevents constitutive heterochromatin and lamina association from spreading into neighboring facultative heterochromatic regions.^51^

**Figure 5.**
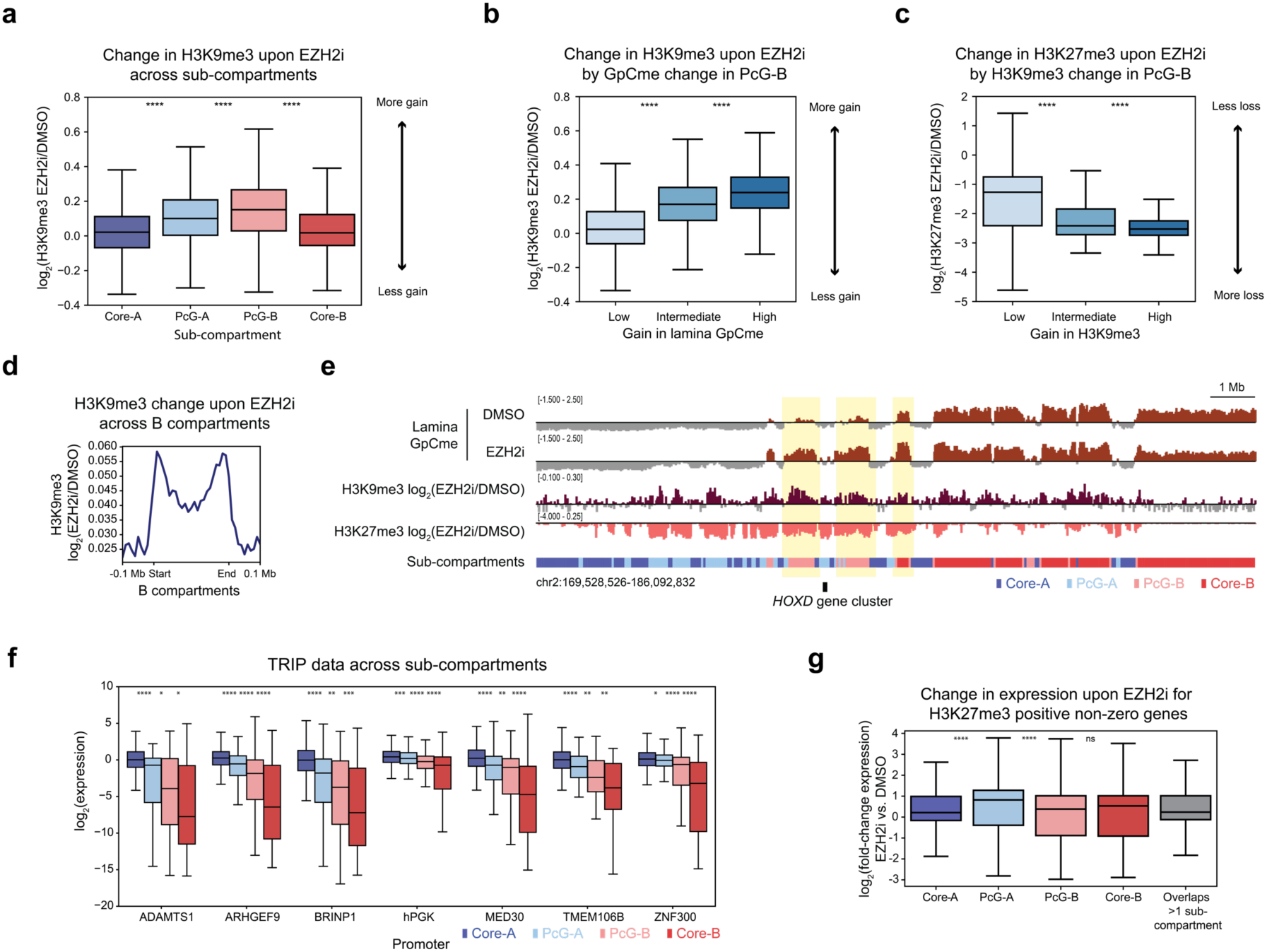
H3K27me3 antagonizes constitutive heterochromatin spreading a) Boxplot showing log_2_ fold-change H3K9me3 ChIP-seq signal between EZH2 inhibition and vehicle treatment for 50 kb bins (*y*-axis) across sub-compartments (*x*-axis). Outlier points are excluded. Significance markers calculated by a Mann-Whitney-Wilcoxon two-sided test are as follows: ns: not significant; *: 0.01 < p ≤ 0.05; **: 0.001 < p ≤ 0.01; ***: 0.0001 < p ≤ 0.001; ****: p ≤ 0.0001. b) Boxplot showing log_2_ fold-change in H3K9me3 levels between EZH2i and vehicle treatment for 50 kb bins (*y*-axis) segregated into three equally sized quantiles by change in *z*-score normalized lamina GpC methylation (*x*-axis) for PcG-B domains. Outlier points are excluded. Significance markers calculated by a Mann-Whitney-Wilcoxon two-sided test are as follows: ns: not significant; *: 0.01 < p ≤ 0.05; **: 0.001 < p ≤ 0.01; ***: 0.0001 < p ≤ 0.001; ****: p ≤ 0.0001. c) Boxplot showing log_2_ fold-change in H3K27me3 levels between EZH2i and vehicle treatment for 50 kb bins (*y*-axis) segregated into three equally sized quantiles by log_2_ fold-change in H3K9me3 (*x*-axis) for PcG-B domains. Outlier points are excluded. Significance markers calculated by a Mann-Whitney-Wilcoxon two-sided test are as follows: ns: not significant; *: 0.01 < p ≤ 0.05; **: 0.001 < p ≤ 0.01; ***: 0.0001 < p ≤ 0.001; ****: p ≤ 0.0001. d) Aggregate profile plot of log_2_ fold-change H3K9me3 ChIP-seq signal between EZH2 inhibition and vehicle treatment (*y*-axis) across B compartment regions (*x*-axis). e) Genome browser tracks of *z*-score normalized lamina GpC methylation levels averaged across replicates for the LIMe-Hi-C data, log_2_ fold-change H3K9me3 ChIP-seq, and log_2_ fold-change H3K27me3 ChIP-seq signal between EZH2i and vehicle treatment near the *HOXD* locus. f) Boxplot of log_2_(TRIP expression) (*y*-axis) across sub-compartments for promoters (*x*-axis). Outlier points are excluded. Published TRIP data is specified in Supplementary Table 1. Significance markers calculated by a Mann-Whitney-Wilcoxon two-sided test are as follows: ns: not significant; *: 0.01 < p ≤ 0.05; **: 0.001 < p ≤ 0.01; ***: 0.0001 < p ≤ 0.001; ****: p ≤ 0.0001. g) Boxplot of log_2_(fold-change expression) (*y*-axis) between EZH2 inhibition and vehicle treatment for genes that overlap with H3K27me3 peaks across sub-compartments (*x*-axis). Outlier points are excluded. Significance markers calculated by a Mann-Whitney-Wilcoxon two-sided test are as follows: ns: not significant; *: 0.01 < p ≤ 0.05; **: 0.001 < p ≤ 0.01; ***: 0.0001 < p ≤ 0.001; ****: p ≤ 0.0001.

We next considered if heterochromatin segregation impacts transcription. To address this question, we analyzed published K562 data from thousands of reporters integrated in parallel (TRIP),^59^ an assay that assesses how chromatin context influences transcription by measuring the expression of the same reporter transcript along with its random genomic integration position. TRIP shows that PcG-B is on average a less repressive chromatin environment than Core-B, consistent with prior observations that the lamina is a strong mediator of gene silencing (**Fig 5f**). Moreover, PcG-B is more repressive than PcG-A despite PcG-B regions possessing lower average levels of H3K27me3 (**Fig 2c**). These patterns were mostly recapitulated when exclusively considering integrations in H3K27me3 ChIP-seq peaks within PcG-A and PcG-B (**Fig S6c**). These findings further support the notion that Polycomb sub-compartments and Core-B represent distinct forms of heterochromatin that have different capacities to silence gene expression.^52^

Prompted by these observations, we considered if EZH2 inhibition led to sub-compartment-specific changes in gene expression. Upon EZH2 inhibition, we observed upregulation of 301 genes and downregulation of 151 genes (adjusted *p*-value < 0.05) (**Fig S6d**) (**Supplementary Table 5**). H3K27me3-marked genes within PcG-A showed the greatest increases in expression (**Fig 5g, Fig S6e**). By contrast, H3K27me3-marked genes within PcG-B remained largely silenced despite both PcG-A and PcG-B genes being repressed prior to EZH2i treatment (**Fig 5g, Fig S6e-f**). These data indicate that EZH2 inhibition has compartment-specific effects on gene expression, with PcG-A genes poised for reactivation and PcG-B genes remaining inactivated—possibly by the repressive nuclear lamina environment or H3K9me3 spreading (**Fig 6**). Altogether, these results support the model that H3K27me3 insulates facultative heterochromatin from the nuclear lamina and the transcriptionally active portion of the A compartment. More broadly, these findings demonstrate how nuclear localization and compartmentalization modulate Polycomb-mediated gene repression.

**Figure 6.**
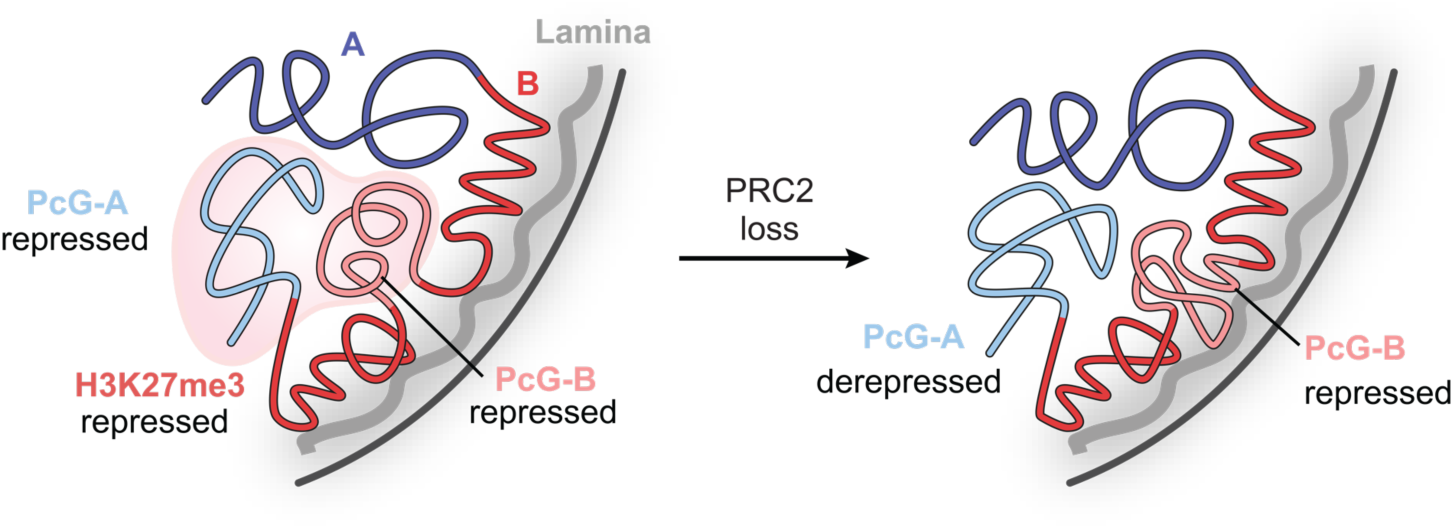
Proposed model of Polycomb-lamina antagonism Schematic of epigenomic changes elicited by loss of PRC2.

## Discussion

In this work, we developed LIMe-Hi-C to simultaneous study lamina association, genome topology, and DNA methylation in a single experiment. The joint profiling of these features enables a global understanding of their interplay and functional consequences, which is often obscured through comparisons of independent datasets.^7^ As we demonstrate, leveraging these multilayered global measurements with chemical inhibition of PRC2 allowed the identification of topologically distinct Polycomb sub-compartments across both the A and B compartments and the direct implication of H3K27me3 in their maintenance.^32,33,35^ These findings are consistent with super-resolution imaging studies demonstrating that Polycomb-rich domains can self-associate and exclude other chromatin states^11^ and previous work partitioning LADs into facultative and constitutive heterochromatin domains.^28,29^ Our work illustrates the multi-faceted impact of H3K27me3 on compartmentalization, complementing previous findings showing the distinct cohesin-independent nature of these interactions.^16,17,37,38,60–62^

Our studies uncovered an antagonism between PRC2 and lamina association, especially for PcG-B regions lying at the borders of LADs and compartments, suggesting a major role of H3K27me3 in segregating heterochromatin subtypes near the nuclear periphery. These findings not only suggest how constitutive and facultative chromatin domains form topologically distinct sub-compartments but also highlight how their modes of lamina contact are independently regulated, with H3K27me3 opposing lamina contact and constitutive heterochromatin self-interaction. These observations were unexpected, as H3K27me3 has been hypothesized to mediate lamina association.^14,25,53^ We propose that this discrepancy might arise from H3K27me3 facilitating peripheral positioning near the lamina, likely at LAD boundaries, but excluding direct contact. This model is consistent with the enrichment of H3K27me3 within the nuclear interior as well as the anti-correlation between lamina association and H3K27me3 levels observed in single cell DamID experiments.^4,63–65^ Furthermore, the antagonism between Polycomb-marked PcG-B and constitutively-marked Core-B regions supports prior findings detailing the differential regulation of facultative and constitutive heterochromatin evidenced upon lamin triple knock-out in mouse embryonic stem cells.^28,29^

The coordinated interplay among H3K27me3, the nuclear lamina, and compartmentalization has broad consequences on Polycomb-mediated repression of gene expression. While H3K27me3-marked genes within PcG-A are activated upon EZH2 inhibition, H3K27me3-marked genes within PcG-B remain largely silenced, consistent with PcG-B repositioning to the nuclear lamina and gaining H3K9me3 to a greater extent. These increases in constitutive heterochromatic features may prevent gene re-activation as both the lamina and HP1*α*, which binds H3K9me3 and interacts with nuclear lamina proteins, are implicated in gene repression.^3,59,66^ Although Polycomb and the nuclear lamina both mediate gene silencing within the B compartment, maintaining two separate forms of heterochromatin may prevent constitutive silencing of Polycomb-genes, enabling their expression in certain contexts (e.g., differentiation).^67,68^ Moreover, the less transcriptionally permissive nature of PcG-B compared to PcG-A reflects how spatial segregation within A compartment regions might poise Polycomb-repressed genes for reactivation. These findings have implications on the functional impact of PRC2 inhibitors, many of which are under clinical evaluation. Altogether, our study illustrates how the complex interplay among Polycomb, A/B compartmentalization, and the nuclear lamina regulates genome structure and gene expression, more broadly highlighting how the cell integrates many regulatory inputs to ensure that gene expression programs are not only stable but also tunable.

## Data availability

Raw and processed LIMe-Hi-C, LIMe-ID, ChIP-seq, and RNA-seq data have been deposited on GEO. Published data analyzed in this study are detailed in Supplementary Table 1. LIMe LAD region calls are supplied in Supplementary Table 2. Sub-compartment calls are supplied in Supplementary Table 3. Regions gaining lamina attachment upon EZH2i are supplied in Supplementary Table 4. Differentially expressed genes are supplied in Supplementary Table 5.

## Code availability

Code employed for analyzing the LIMe-Hi-C and LIMe-ID data is available upon request.

## Acknowledgements

We acknowledge members of the Liau lab for useful discussions. We acknowledge P. van Galen for feedback on the manuscript. We acknowledge K. Zhao, S. Miller, J. Doman, T. Huang, and T. Lu for assistance with high-throughput sequencing. We acknowledge J. Nelson and Z. Niziolek for assistance with cell sorting. We acknowledge the FAS Division of Science Research Computing Group at Harvard University for assistance with cluster computing. A.P.S was supported by the Herchel Smith Graduate fellowship. N.Z.L. was supported by a NSF Graduate Research Fellowship (grant no. DGE1745303). This research was supported by funds from Harvard University and the Broad Institute.

## Author contributions

A.P.S., S.A.R., and B.B.L. conceived the project. A.P.S., S.A.R., E.Z., and B.B.L. developed the LIMe-Hi-C approach. A.P.S. designed, performed, and analyzed the molecular biology experiments. A.P.S. designed, performed, and analyzed the LIMe-Hi-C and LIMe-ID experiments and integrated published data into the study. S.A.R. designed, performed, and analyzed the RNA-seq experiments. A.P.S. and H.R. designed, performed, and analyzed the ChIP-seq experiments. N.Z.L. synthesized the DNMT1 inhibitor. A.P.S. and C.C.W performed the multi-modal analysis of the LIMe-Hi-C data. A.P.S. and B.B.L. wrote the manuscript. S.A.R., N.Z.L., H.R., S.E.J., and M.J.A. edited the manuscript. S.E.J. provided guidance on the experimental strategy and the interpretation of data. M.J.A. designed and oversaw the computational analysis and interpretation of data. B.B.L. designed and oversaw the experimental strategy, computational analysis, and interpretation of data and held overall responsibility for the study.

## Competing interests

B.B.L is on the Scientific Advisory Board of H3 Biomedicine.

**Figure S1.**
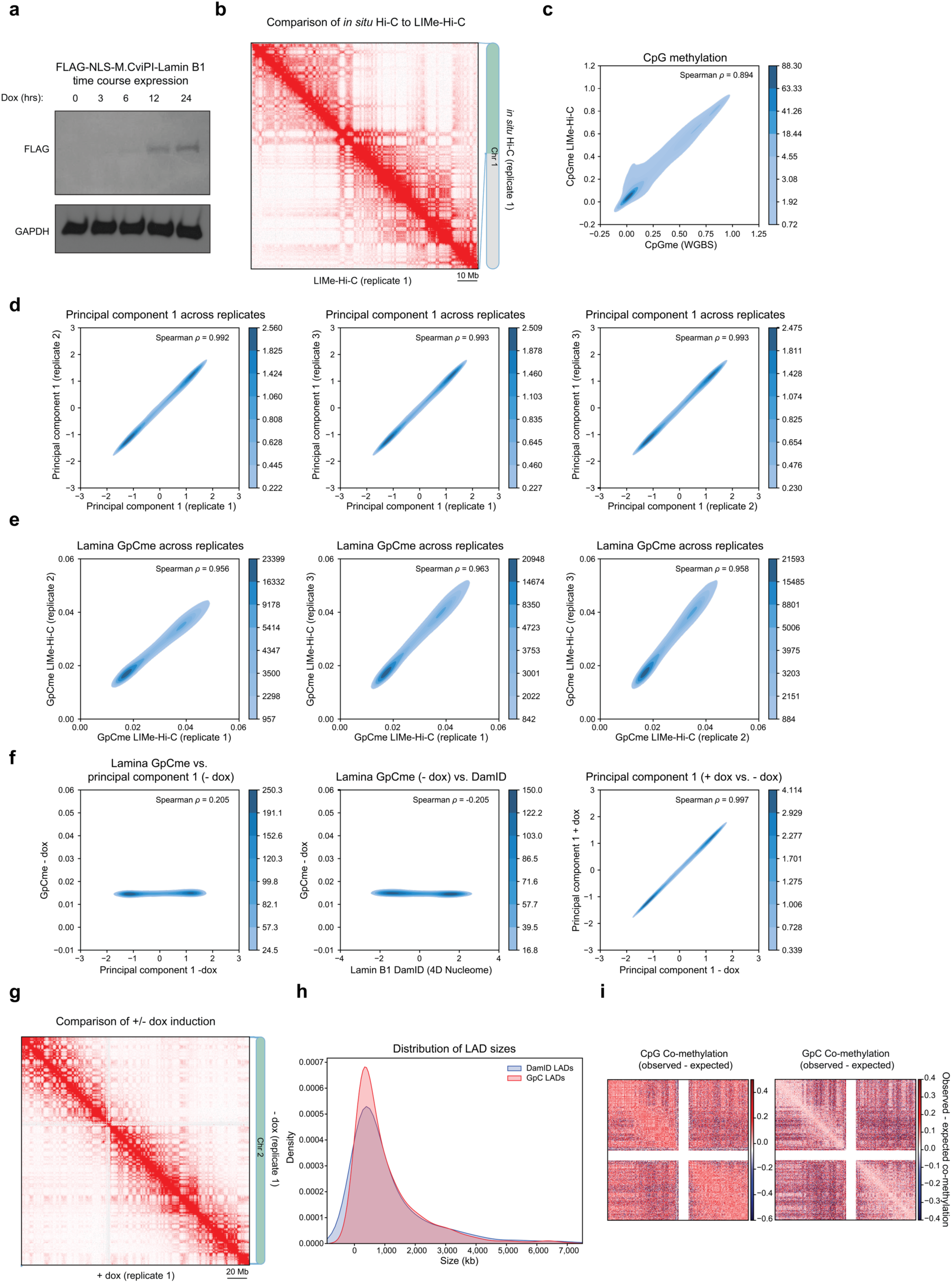
LIMe-Hi-C measurements of LADs, CpG methylation, and Hi-C contacts are robust a) Immunoblot depicting levels of Flag-NLS-M.CviPI-Lamin B1 after specified amount of time of induction with doxycycline. GAPDH was utilized as a loading control. b) Hi-C contact map comparing LIMe-Hi-C (left) to *in situ* Hi-C (right) data for replicate 1 at 250 kb resolution for a region on chromosome 1. c) Density heatmap comparing published whole genome bisulfite sequencing (WGBS) CpG methylation fraction (*x*-axis) to replicate-averaged LIMe-Hi-C CpG methylation fraction (*y*-axis) across 50 kb bins. Published WGBS dataset is specified in Supplementary Table 1. d) Density heatmap comparing principal component 1 in an individual specified replicate (*x*-axis) to principal component 1 in a different specified replicate (*y*-axis) across 50 kb bins for + dox LIMe-Hi-C replicates. e) Density heatmap comparing GpC methylation fraction in an individual specified replicate (*x*-axis) to GpC methylation fraction in a different specified replicate (*y*-axis) across 50 kb bins for + dox LIMe-Hi-C replicates. f) Density heatmap comparing replicate-average GpC methylation fraction (*y*-axis) to replicate-averaged principal component 1 (*x*-axi*s*) across 50 kb bins for the - dox LIMe-Hi-C condition (left). Density heatmap comparing replicate-averaged GpC methylation fraction (*y*-axis) for the - dox LIMe-Hi-C condition to published K562 DamID data (*x*-axis) across 50 kb bins (middle). Density heatmap comparing replicate-averaged principal component 1 across 50 kb bins for the + dox (*y*-axis) and - dox (*x*-axis) LIMe-Hi-C conditions (right). Published DamID dataset is specified in Supplementary Table 1. g) Hi-C contact map comparing + dox (left) and - dox (right) conditions for replicate 1 at 250 kb resolution. h) Density plot depicting the relative density (*y*-axis) of region sizes (*x*-axis) for LIMe and DamID LADs. The density function for each LAD type is independently scaled. i) Observed - expected per-read CpG co-methylation (left) and GpC co-methylation (right) fraction for both arms of chromosome 1 (see Methods). Regions with low coverage near the centromere are marked in white.

**Figure S2.**
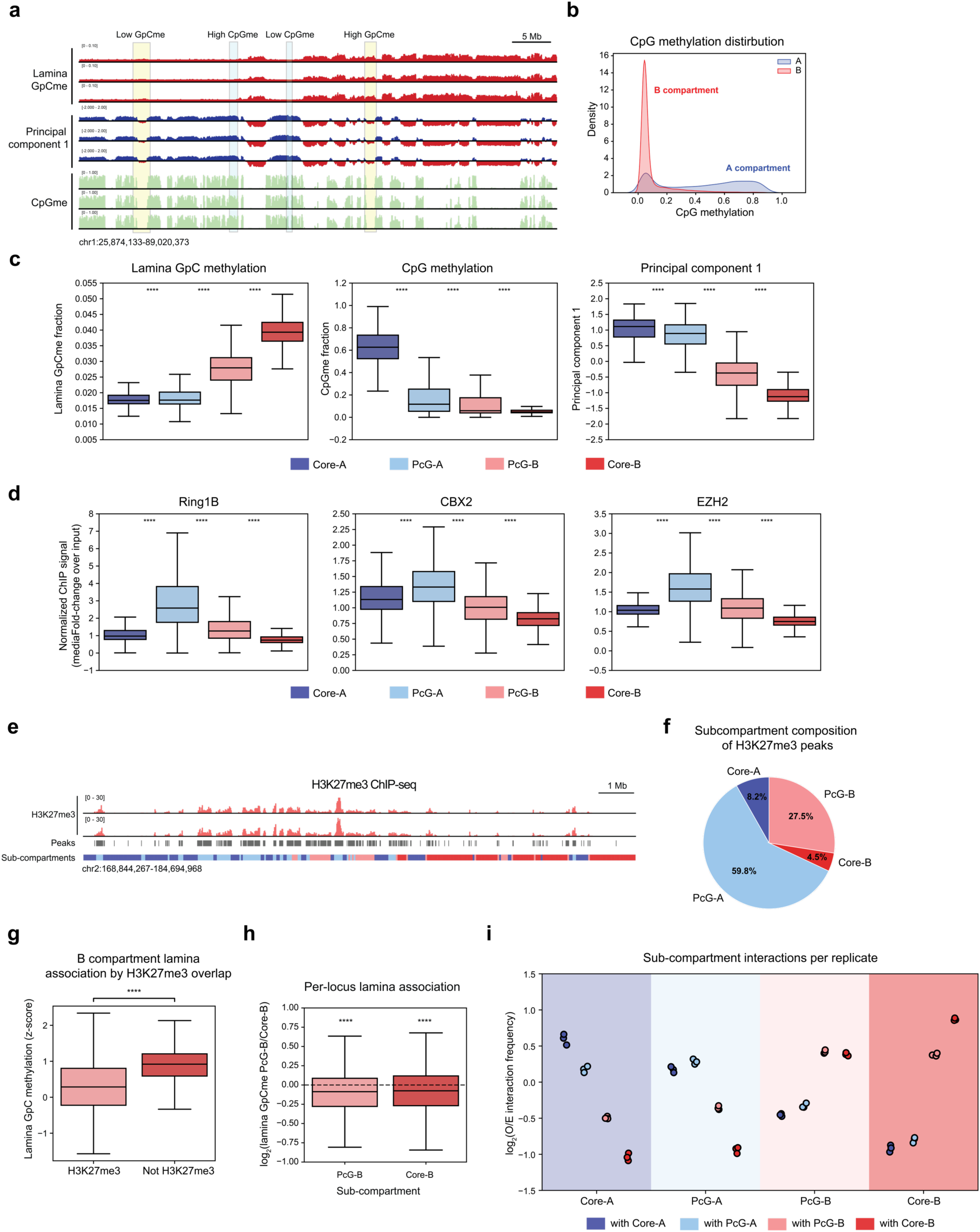
Sub-compartments identified in K562 are epigenomically distinct a) Genome browser tracks of lamina GpC methylation, principal component 1 and CpG methylation in K562 for LIMe-Hi-C replicates depicting regions within the B compartment with variable lamina GpC methylation and regions within the A compartment with variable CpG methylation levels. b) Density plot depicting the relative density (*y*-axis) of fraction open sea CpG methylation (excluding CpG islands) (*x*-axis) for 50 kb bins across A and B compartments in K562. The density function for each compartment is independently scaled. c) Boxplot of LIMe-Hi-C features (*y*-axis: fraction GpC methylation (left), fraction CpG methylation (middle), and principal component 1 (right)) across the sub-compartments (*x*-axis) in K562. Outlier points are excluded. Significance markers calculated by a Mann-Whitney-Wilcoxon two-sided test are as follows: ns: not significant; *: 0.01 < p ≤ 0.05; **: 0.001 < p ≤ 0.01; ***: 0.0001 < p ≤ 0.001; ****: p ≤ 0.0001. d) Boxplot showing levels of Polycomb factors (*y*-axis, fold-change over input/median signal) for 50 kb bins across sub-compartments (*x*-axis) in K562. Outlier points are excluded. Published datasets are specified in Supplementary Table 1. Significance markers calculated by a Mann-Whitney-Wilcoxon two-sided test are as follows: ns: not significant; *: 0.01 < p ≤ 0.05; **: 0.001 < p ≤ 0.01; ***: 0.0001 < p ≤ 0.001; ****: p ≤ 0.0001. Published ChIP-seq datasets are specified in Supplementary Table 1. e) Genome browser tracks of H3K27me3 ChIP-seq signal with H3K27me3 peaks and sub-compartment designations below. f) Pie chart depicting fraction base pair overlap of the different sub-compartments with H3K27me3 peaks. g) Boxplot of *z*-score normalized lamina GpC methylation (*y*-axis) for 50 kb bins within the B compartment by their overlap with H3K27me3 peaks (*x*-axis). Outlier points are excluded. Significance markers calculated by a Mann-Whitney-Wilcoxon two-sided test are as follows: ns: not significant; *: 0.01 < p ≤ 0.05; **: 0.001 < p ≤ 0.01; ***: 0.0001 < p ≤ 0.001; ****: p ≤ 0.0001. h) Boxplot for all 50 kb bins on chromosome 1 across PcG-B and Core-B (*x*-axis) depicting the log_2_ ratio of the interval’s lamina association status (*y*-axis) if it is interacting with the PcG-B versus Core-B. Outlier points are excluded. Significance markers were calculated by a one sample one-sided t-test to determine if the mean of the population was less than 0 and are represented as follows: ns: not significant; *: 0.01 < p ≤ 0.05; **: 0.001 < p ≤ 0.01; ***: 0.0001 < p ≤ 0.001; ****: p ≤ 0.0001. i) Dotplot showing log_2_(observed/expected contact frequency) for every replicate, averaged across the genome (*y*-axis) among all sub-compartments (*x*-axis) for K562. O/E denotes observed/expected.

**Figure S3.**
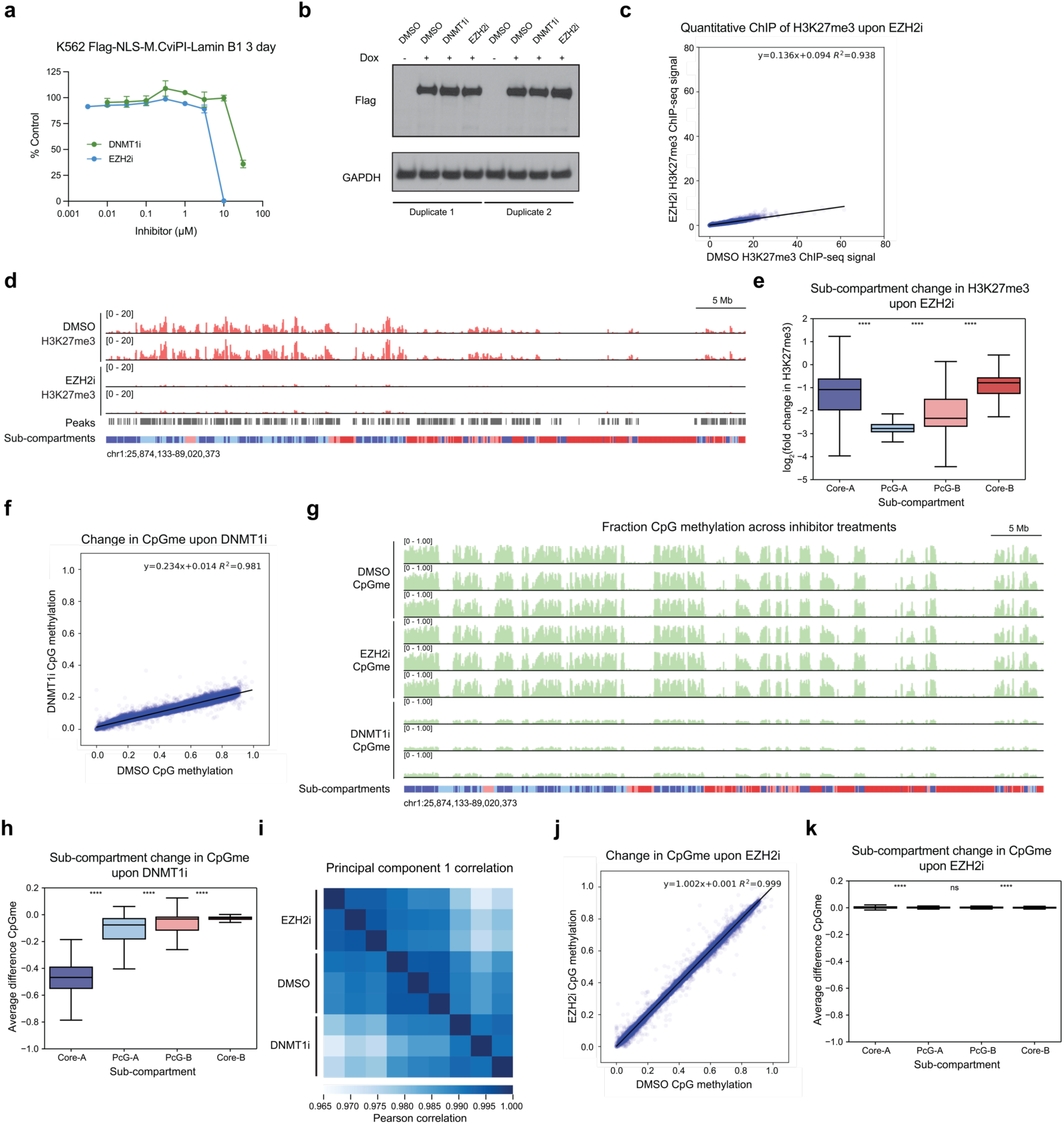
Inhibition of EZH2 depletes H3K27me3 and inhibition of DNMT1 depletes DNA methylation globally a) Dose-response curve of concentration inhibitor (*x*-axis) and percent growth relative to vehicle control (*y*-axis) for Flag-NLS-M.CviPI-Lamin B1 cells treated with DNMT1i and EZH2i. Data represent mean ± s.e.m across three technical replicates with the experiment performed once. b) Immunoblot for the Flag-NLS-M.CviPI-Lamin B1 fusion protein following treatment with EZH2i, DNMT1i, or DMSO with and without doxycycline. GAPDH was utilized as a loading control. The experiment was performed once in duplicate. c) Scatterplot comparing replicate-averaged H3K27me3 quantitative ChIP-seq levels for DMSO (*x*-axis) and EZH2 inhibitor (*y*-axis) treatments for 50 kb bins. Equation for the line of best fit is depicted in the plot. d) Genome browser tracks of H3K27me3 quantitative ChIP-seq signal for EZH2i and vehicle treatments for both replicates. e) Boxplot showing log_2_ fold-change H3K27me3 ChIP-seq signal between EZH2 inhibition and vehicle treatment for 50 kb bins (*y*-axis) across sub-compartments (*x*-axis). Outlier points are excluded. Significance markers calculated by a Mann-Whitney-Wilcoxon two-sided test are as follows: ns: not significant; *: 0.01 < p ≤ 0.05; **: 0.001 < p ≤ 0.01; ***: 0.0001 < p ≤ 0.001; ****: p ≤ 0.0001. f) Scatterplot comparing replicate-averaged CpG methylation fraction between DMSO (*x*-axis) and DNMT1 inhibitor (*y*-axis) treatments for 50 kb bins. Equation for the line of best fit is depicted in the plot. g) Genome browser tracks of the CpG methylation fraction across EZH2i, DNMT1i, and vehicle treatments for all three replicates. h) Boxplot showing difference in CpG methylation fraction between DNMT1 inhibition and vehicle treatment for 50 kb bins (*y*-axis) across sub-compartments (*x*-axis). Outlier points are excluded. Significance markers calculated by a Mann-Whitney-Wilcoxon two-sided test are as follows: ns: not significant; *: 0.01 < p ≤ 0.05; **: 0.001 < p ≤ 0.01; ***: 0.0001 < p ≤ 0.001; ****: p ≤ 0.0001. i) Pearson correlation heatmap of principal component 1 across 50 kb bins for LIMe-Hi-C data across replicates and inhibitor treatment conditions. j) Scatterplot comparing replicate-averaged CpG methylation fraction between DMSO (*x*-axis) and EZH2 inhibitor (*y*-axis) treatments for 50 kb bins. Equation for the line of best fit is depicted in the plot. k) Boxplot showing difference in CpG methylation fraction between EZH2 inhibition and vehicle treatment for 50 kb bins (*y*-axis) across sub-compartments (*x*-axis). Outlier points are excluded. Significance markers calculated by a Mann-Whitney-Wilcoxon two-sided test are as follows: ns: not significant; *: 0.01 < p ≤ 0.05; **: 0.001 < p ≤ 0.01; ***: 0.0001 < p ≤ 0.001; ****: p ≤ 0.0001.

**Figure S4.**
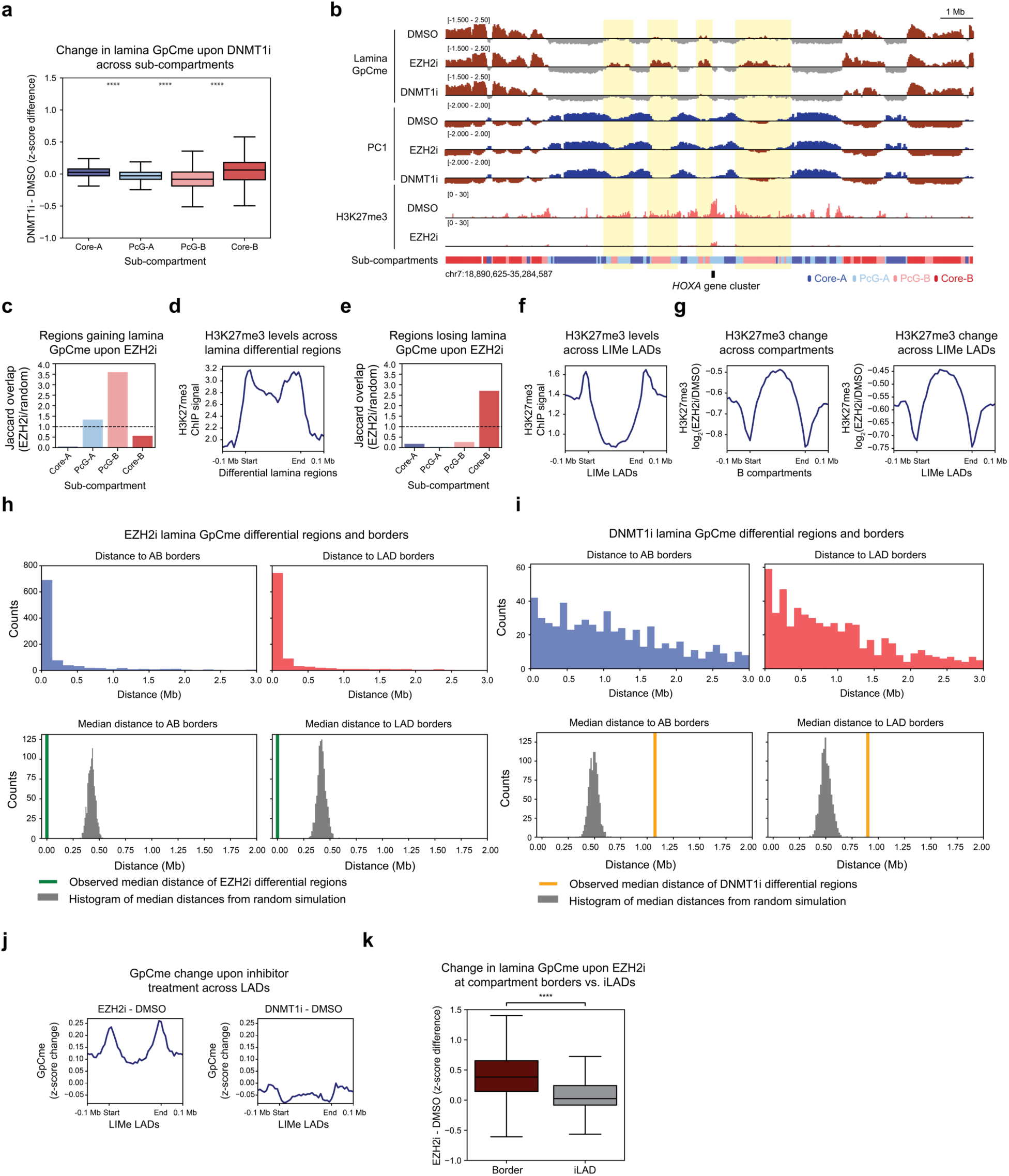
Regions gaining lamina association upon EZH2 inhibition are enriched within PcG-B and reside at LAD and compartment borders a) Boxplot showing the difference in *z*-score normalized GpC methylation between DNMT1i and vehicle treatment for 50 kb bins (*y*-axis) across sub-compartments (*x*-axis). Outlier points are excluded. Significance markers calculated by a Mann-Whitney-Wilcoxon two-sided test are as follows: ns: not significant; *: 0.01 < p ≤ 0.05; **: 0.001 < p ≤ 0.01; ***: 0.0001 < p ≤ 0.001; ****: p ≤ 0.0001. b) Genome browser tracks of *z*-score normalized lamina GpC methylation levels, principal component 1, and H3K27me3 ChIP-seq signal averaged across the replicates for LIMe-Hi-C and quantitative ChIP-seq data near the *HOXA* locus. c) Barplot across all sub-compartments (*x*-axis) showing the ratio of the Jaccard overlap for regions gaining lamina contact upon EZH2 inhibition with the specified sub-compartment relative to a simulated random distribution (*y*-axis, observed Jaccard overlap/mean Jaccard overlap from simulation). d) Aggregate profile plot of baseline H3K27me3 ChIP-seq signal for vehicle treatment (*y*-axis) across regions gaining lamina contact upon EZH2 inhibition (*x*-axis) (see Methods). e) Barplot across all sub-compartments (*x*-axis) showing the ratio of the Jaccard overlap for regions losing lamina contact upon EZH2 inhibition with the specified sub-compartment relative to a simulated random distribution (*y*-axis, observed Jaccard overlap/mean Jaccard overlap from simulation). f) Aggregate profile plot of baseline H3K27me3 ChIP-seq signal for vehicle treatment (*y*-axis) across LIMe LADs (*x-axis*). g) Aggregate profile plot of log_2_ fold-change H3K27me3 ChIP-seq signal between EZH2 inhibition and vehicle treatment (*y*-axis) across B compartments (left, *x*-axis) and LIMe LADs (right, *x*-axis). h) Histogram of the frequency (*y*-axis) of genomic distances (*x*-axis) for regions gaining lamina contact upon EZH2 inhibition to A/B and LAD borders (top). Histogram of median genomic distances (*x*-axis) of randomly shuffled regions to A/B and LAD borders relative to observed counts (*y*-axis) for a simulated random distance distribution (bottom). i) Histogram of the frequency (*y*-axis) of genomic distances (*x*-axis) for regions gaining lamina contact upon DNMT1 inhibition to A/B and LAD borders (top). Histogram of median genomic distances (*x*-axis) of randomly shuffled regions to A/B and LAD borders relative to observed counts (*y*-axis) for a simulated random distance distribution (bottom). j) Aggregate profile plots depicting change in *z*-score normalized lamina GpC methylation (*y*-axis) across LIMe LADs (*x*-axis) for inhibitor treatments relative to vehicle treatment. k) Boxplot showing the difference in *z*-score normalized GpC methylation between EZH2i and vehicle treatment (*y*-axis) for H3K27me3 peaks within 50 kb of a compartment border versus those peaks greater than 50 kb from a border within the A compartment (*x*-axis). Outlier points are excluded. Significance markers calculated by a Mann-Whitney-Wilcoxon two-sided test are as follows: ns: not significant; *: 0.01 < p ≤ 0.05; **: 0.001 < p ≤ 0.01; ***: 0.0001 < p ≤ 0.001; ****: p ≤ 0.0001.

**Figure S5.**
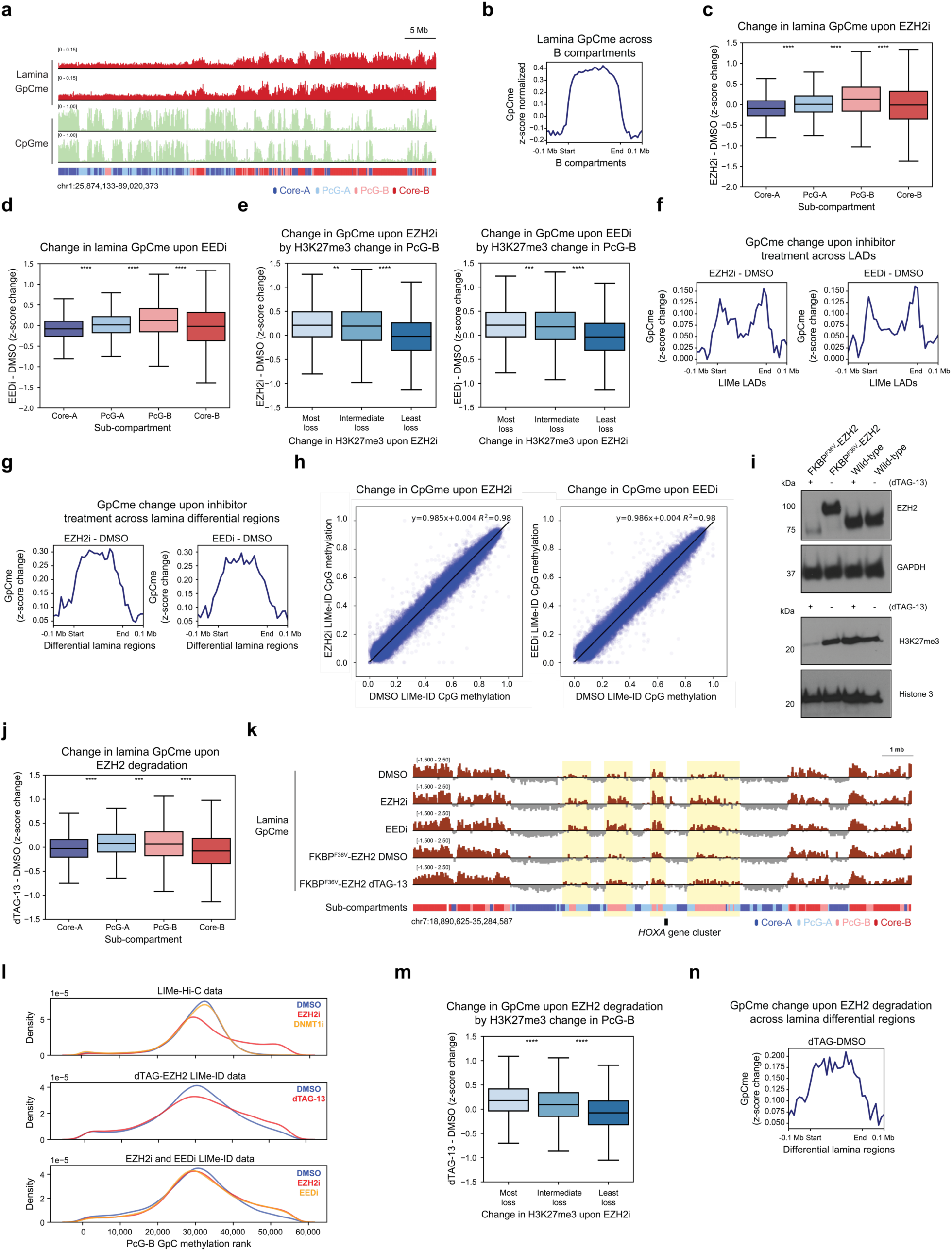
LIMe-ID following PRC2 perturbations support the role of H3K27me3 in antagonizing lamina association a) Genome browser tracks of GpC and CpG methylation fraction for vehicle treatment for both LIMe-ID duplicates with sub-compartment designation below. b) Aggregate profile plot of LIMe-ID replicate-averaged *z*-score normalized GpC methylation (*y*-axis) across B compartments (*x*-axis). c) Boxplot showing the difference in *z*-score normalized GpC methylation between inhibitor and vehicle treatment for 50 kb bins (*y*-axis) across sub-compartments (*x*-axis) for the EZH2 inhibitor LIMe-ID data. Outlier points are excluded. Significance markers calculated by a Mann-Whitney-Wilcoxon two-sided test are as follows: ns: not significant; *: 0.01 < p ≤ 0.05; **: 0.001 < p ≤ 0.01; ***: 0.0001 < p ≤ 0.001; ****: p ≤ 0.0001. d) Boxplot showing the difference in *z*-score normalized GpC methylation between inhibitor and vehicle treatment for 50 kb bins (*y*-axis) across sub-compartments (*x*-axis) for the EED inhibitor LIMe-ID data. Outlier points are excluded. Significance markers calculated by a Mann-Whitney-Wilcoxon two-sided test are as follows: ns: not significant; *: 0.01 < p ≤ 0.05; **: 0.001 < p ≤ 0.01; ***: 0.0001 < p ≤ 0.001; ****: p ≤ 0.0001. e) Boxplot showing the difference in *z*-score normalized GpC methylation between inhibitor and vehicle treatment for 50 kb bins (*y*-axis) segregated into three equally sized quantiles by log_2_ fold-change in H3K27me3 (*x*-axis) for PcG-B domains for both EZH2 inhibitor LIMe-ID data (left) and EED inhibitor LIMe-ID data (right). Outlier points are excluded. Significance markers calculated by a Mann-Whitney-Wilcoxon two-sided test are as follows: ns: not significant; *: 0.01 < p ≤ 0.05; **: 0.001 < p ≤ 0.01; ***: 0.0001 < p ≤ 0.001; ****: p ≤ 0.0001. f) Aggregate profile plots depicting change in *z*-score normalized lamina GpC methylation (*y*-axis) across LIMe LADs (*x*-axis) upon EZH2 inhibition (left) and EED inhibition (right) (see Methods). g) Aggregate profile plots depicting change in *z*-score normalized lamina GpC methylation (*y*-axis) across EZH2 LIMe-Hi-C lamina differential regions (*x*-axis) for EZH2 inhibition (left) and EED inhibition (right) (see Methods). h) Scatterplot comparing CpG methylation fraction between DMSO (*x*-axis) and EZH2 inhibitor treatment (left, *y*-axis) and EED inhibitor treatment (right, *y*-axis) for 50 kb bins averaged across replicates. Equation for the line of best fit is depicted in the plot. i) Immunoblot depicting levels of EZH2 for the N-terminal EZH2 knock-in cell line and wild-type K562 upon DMSO and dTAG-13 treatment. GAPDH was utilized as a loading control (top). Histone immunoblot depicting levels of H3K27me3 for the N-terminal EZH2 knock-in cell line and wild-type K562 upon DMSO and dTAG-13 treatment. Histone 3 was utilized as a loading control (bottom). j) Boxplot showing the difference in *z*-score normalized GpC methylation between dTAG-13 and vehicle treatment for 50 kb bins (*y*-axis) across sub-compartments (*x*-axis). Outlier points are excluded. Significance markers calculated by a Mann-Whitney-Wilcoxon two-sided test are as follows: ns: not significant; *: 0.01 < p ≤ 0.05; **: 0.001 < p ≤ 0.01; ***: 0.0001 < p ≤ 0.001; ****: p ≤ 0.0001. k) Genome browser tracks of *z*-score normalized lamina GpC methylation levels averaged across replicates for LIMe-ID inhibitor treatments and EZH2 degradation near the *HOXA* locus. l) Density plot of GpC methylation rank for PcG-B regions for the LIMe-Hi-C inhibitor treatments as well as the LIMe-ID inhibitor treatments and EZH2 degradation data. m) Boxplot showing the difference in *z*-score normalized GpC methylation between dTAG-13 and vehicle treatment for 50 kb bins (*y*-axis) segregated into three equally sized quantiles by log_2_ fold-change in H3K27me3 (*x*-axis) for PcG-B. Outlier points are excluded. Significance markers calculated by a Mann-Whitney-Wilcoxon two-sided test are as follows: ns: not significant; *: 0.01 < p ≤ 0.05; **: 0.001 < p ≤ 0.01; ***: 0.0001 < p ≤ 0.001; ****: p ≤ 0.0001. n) Aggregate profile plot depicting change in *z*-score normalized lamina GpC methylation (*y*-axis) across EZH2 LIMe-Hi-C lamina differential regions (*x*-axis) for dTAG-13 treatment (see Methods).

**Figure S6.**
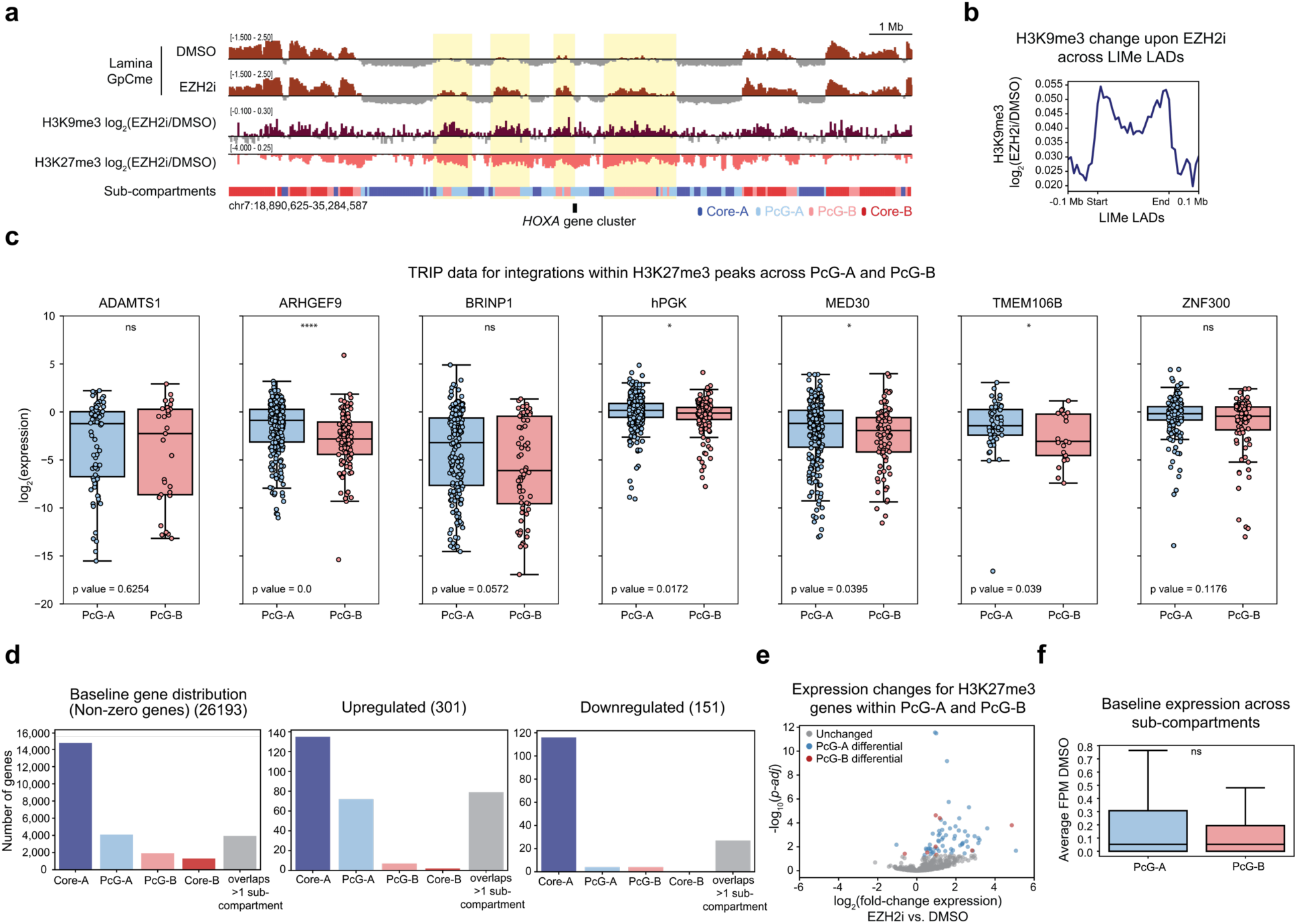
Polycomb sub-compartments represent distinct transcriptional environments a) Genome browser tracks of *z*-score normalized lamina GpC methylation levels averaged across replicates for the LIMe-Hi-C data, log_2_ fold-change H3K9me3 ChIP-seq, and log_2_ fold-change H3K27me3 ChIP-seq signal between EZH2i and vehicle treatment near the *HOXA* locus. b) Aggregate profile plot of log_2_ fold-change H3K9me3 ChIP-seq signal between EZH2 inhibition and vehicle treatment (*y*-axis) across LIMe LADs (*x*-axis). c) Boxplot with individual points overlaid of log_2_(TRIP expression) (*y*-axis) across the PcG-A and PcG-B sub-compartments (*x*-axis) for promoter integrations that overlap with H3K27me3 peaks. Significance markers calculated by a Mann-Whitney-Wilcoxon two-sided test are as follows: ns: not significant; *: 0.01 < p ≤ 0.05; **: 0.001 < p ≤ 0.01; ***: 0.0001 < p ≤ 0.001; ****: p ≤ 0.0001. d) Bar charts depicting number of genes within each sub-compartment for those that are detected by RNA-seq (non-zero) and those that are either upregulated or downregulated upon EZH2 inhibition. e) Volcano plot depicting -log_10_(adjusted *p*-value) (*y*-axis) relative to log_2_(fold-change expression) (*x*-axis) between EZH2 inhibition and vehicle treatment for H3K27me3-positive PcG-A and PcG-B genes. f) Boxplot of baseline expression (*y*-axis, fragment per million mapped fragments (FPM)) for vehicle treatment of non-zero genes that overlap with H3K27me3 peaks across PcG-A and PcG-B (*x*-axis). Outlier points are excluded. Significance markers calculated by a Mann-Whitney-Wilcoxon two-sided test are as follows: ns: not significant; *: 0.01 < p ≤ 0.05; **: 0.001 < p ≤ 0.01; ***: 0.0001 < p ≤ 0.001; ****: p ≤ 0.0001.

## Methods

### Cell culture and lentivirus production

K562 was obtained from ATCC. HEK 293T was obtained from Life Technologies. K562 was authenticated by Short Tandem Repeat profiling (Genetica) and routinely tested for mycoplasma (Sigma-Aldrich). All cell lines were cultured in a humidified 5% CO_2_ incubator at 37 °C with media supplemented with 100 U/ml penicillin and 100 µg/ml streptomycin (Life Technologies). K562 was cultured in RPMI-1640 (Life Technologies) with 10% fetal bovine serum (FBS) (Peak Serum). HEK 293T was cultured in DMEM (Life Technologies) with 10% FBS (Peak Serum). For lentivirus production, expression plasmids were co-transfected with *GAG/POL* and *VSVG* plasmids into HEK 293T cells using FuGENE HD (Promega). Media was exchanged after 6-10 h, and the viral supernatant was collected 48-72 h after transfection and filtered (0.45 µm). K562 was transduced by spinfection at 2,000*g* for 90 min at 37 °C with 8 µg/ml polybrene (Santa Cruz Biotechnology). To generate the M.CviPI overexpression bulk cell lines, 48 h post-transduction, geneticin (Life Technologies) selection was carried out for approximately one week at 0.6 mg/ml. Single cell clones were isolated from the bulk cell line through fluorescence-activated cell sorting, and the expression of the fusion was validated through immunoblot.

### Mammalian overexpression constructs

The Lamin B1 open reading frame (ORF) was subcloned from mCherry-LaminB1-10, a gift from M. Davidson (Addgene #55609), and the M.CviPI ORF with a C-terminal glycine-serine (GS) linker (GGSGGGS) was ordered as a gene block (Integrated DNA Technologies). An N-terminal FLAG tag and a nuclear localization signal (NLS) along with the M.CvIPI ORF were then cloned N-terminal to Lamin B1 into pENTR (Invitrogen) using Gibson cloning (New England Biolabs). The final overexpression construct for FLAG-NLS-M.CviPI-LaminB1 in pInducer20 (Invitrogen) was obtained through a recombination reaction using Gateway cloning (LR Clonase™ II, Thermo Fisher) between the M.CviPI-pENTR construct and pInducer20. For the LIMe-ID experiments with EED and EZH2 inhibition, the M.CviPI-LaminB1 plasmid without the FLAG-NLS sequence was employed instead and cloned in an analogous manner.

### Generating dTAG-EZH2 knock-in

EZH2 homology arms of 1 kb were amplified from K562 genomic DNA and cloned into the ph1 backbone along with Puro-P2A-dTAG, sub-cloned from pCRIS-PITChv2-Puro-dTAG, a gift from J. Bradner (Addgene # 91793). The sgRNA, 5’-GAGAAGGGACCAGTTTGTTGG-3’ that was previously validated for N-terminal EZH2 knock-in^69^ was cloned into PX459 by restriction enzyme digestion. Wild type K562 cells were transfected with both the donor and guide plasmids by nucleofection (Lonza, program T016). Cells were selected 3 days after transfection with puromycin (2µg/mL) (Thermo Fisher Scientific) for a duration of 5 days. Single cell clones were isolated from the bulk cell line through fluorescence-activated cell sorting and validated by western blot and sanger sequencing. The Flag-NLS-M.CvIPI-LaminB1 construct was then transduced into this clonal knock-in cell line and selection was carried out with geneticin as specified above. The cell line resulting from this transduction was then used directly for subsequent LIMe-ID experiments.

### Immunoblotting

Fusion protein overexpression was induced by treatment with doxycycline (dox) (Sigma Aldrich) at a concentration of 1µg/mL for the specified amount of time. Pellets were harvested for western blot analysis and washed with PBS (Corning). Cells were subsequently lysed on ice using RIPA buffer (Boston BioProducts) supplemented with fresh HALT Protease Inhibitor (Thermo Fisher Scientific) and EDTA (Thermo Fisher Scientific), and the lysates were clarified through centrifugation. For histone immunoblotting, histones were extracted according to the recommended manufacturer’s one-step protocol using the Histone Extraction Kit (Active Motif). The protein concentration of the lysates was determined using the BCA Protein Assay Kit (Thermo Fisher Scientific). Immunoblotting was performed according to standard procedures. To assess histone loading, the blot was stripped with Restore Western Blot Stripping Buffer (Thermo Fisher Scientific) for 30 min prior to re-probing for histone H3. The primary antibodies used for immunoblotting are as follows: GAPDH (Santa Cruz Biotechnology sc-477724); monoclonal anti-FLAG M2 (Sigma-Aldrich F1804); Tri-Methyl-Histone H3 Lys27 (Cell Signaling Technology C36B11); Histone H3 (Cell Signaling Technology 9715S) and EZH2 (Cell Signaling Technology 5246).

### Cell growth assays

Flag-NLS-M.CviPI-Lamin B1 K562 cells were plated in triplicate at a density of 4000/well in a 96-well plate. Drug or vehicle (DMSO) were dosed at the specified concentration for 3 days. After 3 days of treatment, cell viability was measured using CellTiter-Glo (Promega) with the luminescence detector on the SpectraMax i3x plate reader, and data were processed using Prism.

### Inhibitor treatment for Hi-C and RNA-seq

The Flag-NLS-M.CviPI-LaminB1 clonal cell line was seeded at a density of 125K/mL and treated with 10 µM GSK3482364 (synthesized as previously described)^48^, 1 µM GSK343 (Sigma Aldrich) or 0.1% DMSO (Sigma Aldrich) for a total of three days. Two days after the start of the inhibitor or vehicle treatment, doxycycline (Sigma Aldrich) was added to the cells at a final concentration of 1 µg/mL for the remaining 24 hours of the drug treatments. For the no induction condition, blank media was added in place of doxycycline. Upon completion of the drug treatments, pellets were taken for LIMe-Hi-C and RNA-seq in triplicate.

### Hi-C and LIMe-Hi-C

Cells were treated in triplicate with either GSK343, GSK3482364, or vehicle and induced with doxycycline as specified above. *In situ* Hi-C was performed according to a published protocol.^2^ Briefly, 4 × 10^6^ cells were fixed at a density of 10^6^ cell/mL in media containing 10% FBS and 1% formaldehyde for 10 min with rotation. Formaldehyde was quenched with glycine (final concentration, 0.2 M), and cell pellets were washed with PBS, and flash frozen. Nuclei were lysed, and overnight digestion was performed using 200 U of DpnII at 37 °C with shaking (New England Biolabs, R0543L) in NEB buffer 3.1. Heat inactivation was performed by incubating the samples at 65 °C for 20 min. Digested ends were filled in with biotinylated dATP (Life Technologies, 19524-016) for 30 min at 37 °C, and *in situ* ligation was performed with T4 DNA ligase (New England Biolabs, M0202L) for 6 hours at room temperature with rotation. Samples were digested with Proteinase K (Sigma Aldrich) at 55 °C for 30 min and heated overnight at 68 °C to reverse the crosslinks. DNA was purified through an ethanol precipitation and sheared to a size of 300-500 bp with sonication using the Covaris sonicator (S220). DNA was size-selected following shearing using a 0.55x followed by a 0.70x double-SPRI magnetic bead purification (Omega Bio-Tek, M1378-01). At this point, the DNA was split in half for bisulfite (LIMe) and traditional *in situ* Hi-C, if performed. For the traditional *in situ* Hi-C workflow, biotinylated fragments were isolated using T1 streptavidin beads (Life Technologies, 65601), and library preparation was performed on beads according to published precedent.^2^ The preparation of bisulfite (LIMe) Hi-C libraries was performed according to the Hi-Culfite protocol.^40^ Specifically, 0.5-1µg of DNA was bisulfite-converted with the EpiTect Fast Bisulfite Conversion Kit (Qiagen, 59824) using two columns per sample according to the manufacturer’s protocol with the following modifications: the two 60 °C bisulfite conversion incubations were extended from 10 min to 20 min each, and DNA was purified without carrier RNA. Following bisulfite conversion, the samples were eluted in final volume 20 µL, and 15 µL of material was used. Biotinylated fragments were purified using C1 streptavidin beads (Life Technologies, 65001). Specifically, 15 µL of streptavidin C1 beads were added to each sample, and the biotinylated DNA was allowed to incubate for 10 min with the beads at 55 °C in 1x denaturing buffer (10 mM Tris pH 7.5, 5 mM EDTA, 500 mM LiCl, 0.5% Igepal CA630, 0.2% SDS, 4 M urea). Following the incubation, the streptavidin beads were washed twice with prewarmed denaturing buffer at 55 °C for 2 min and once with 10 mM Tris-HCl pH 7.5. DNA was eluted in 15 µL of Tris pH 7.5 for 5 min at 95 °C and subsequently quantified using ssDNA Qubit quantification. LIMe-Hi-C libraries were then generated from 25-50 ng of input DNA using Accel-NGS-methyl-Seq Kit (Swift Biosciences) according to the manufacturers protocol with 9 cycles of amplification. Care was taken to avoid freeze-thaw cycles once the DNA was bisulfite-converted prior to library preparation. Quality of the libraries were assessed using a MiSeq Genome Analyzer (Illumina) with 75 bp paired-end reads. Libraries were subsequently sequenced on a NovaSeq S4 kit (Illumina) with 150 bp paired-end reads to a sequencing depth of around 600 million reads per sample.

### Quantitative ChIP-seq

ChIP was performed as previously described in duplicate for Flag-NLS-M.CvIPI-Lamin B1 cells treated with DMSO or GSK343 and induced with doxycycline as described above,^70^ with the following modifications: spike-in Drosophila S2 chromatin (Active Motif, 08221011) at a concentration of 10 ng/million cells and spike-in antibody (Active Motif, 61686) at a concentration of 0.4 µg/million cells were added following lysis and sonication but prior to immunoprecipitation. H3K27me3 immunoprecipitation was performed on 4 million cells with 10 µL of H3K27me3 antibody (Cell Signaling, C36B11 Lot #19). H3K9me3 immunoprecipitation was performed on 3 million cells with 8 µL of H3K9me3 antibody (Cell Signaling, 13969S Lot #3). Samples were sequenced using a NovaSeq SP kit (Illumina) with 50 bp paired-end reads.

### RNA-seq

In triplicate separate cultures, cells were treated with GSK343 or vehicle and induced with doxycycline as described above. Total RNA was isolated using the RNeasy Plus Mini Kit (Qiagen). Library preparation was performed using the Quantseq 3’ mRNA-Seq Library Prep Kit FWD (Lucigen) according to the manufacturer’s protocol. The samples were sequenced with a NovaSeq SP kit (Illumina) for 50 bp single-end reads.

### LIMe-ID

For inhibitor LIMe-ID experiments, M.CvIPI-Lamin-B1 cells were treated in duplicate with 1 µM GSK343 (Selleck Chemicals), 5 µM EED226 (Selleck Chemicals), or vehicle (0.1% DMSO (Sigma Aldrich)) for a total of 3 days and induced with doxycycline as specified above. For degradation LIMe-ID experiments, Flag-NLS-M.CvIPI-Lamin-B1 dTAG-EZH2 knock-in cells were treated in duplicate with either 500 nM dTAG-13 (Sigma Aldrich) or vehicle (0.1% DMSO (Sigma Aldrich)) for a total of 3 days and induced with doxycycline as specified above. Cell pellets of 1 million cells each were harvested by centrifugation and washed with PBS (Corning). DNA was purified through QIAamp DNA blood mini kit (Qiagen) with the addition of RNAse and sheared to a size of 400 bp in 50 µL by sonication using the Covaris sonicator (S220). DNA was subsequently bisulfite converted with the EpiTect Fast Bisulfite Conversion Kit (Qiagen) according to the manufacturer’s protocol with the following modifications: the two 60 °C bisulfite conversion incubations were extended from 10 min to 20 min each, and DNA was purified without carrier RNA. DNA was quantified and libraries were then generated from 25-50 ng of input DNA using Accel-NGS-methyl-Seq Kit (Swift Biosciences) according to the manufacturers protocol with 9 cycles of amplification. Samples were sequenced using a NovaSeq SP kit (Illumina) with 50 bp paired-end reads.

### Hi-C and LIMe-Hi-C data processing

*In situ* Hi-C data were processed according to the SLURM version of Juicer pipeline (version 1.6) with Juicertools (version 1.22.01).^71^ The LIMe-Hi-C data were processed utilizing the CPU version of the JuiceMe (version 1.0.0) pipeline developed for analyzing bisulfite Hi-C data using Juicertools (version 1.22.01) with the following modifications.^40,71^ Reads were aligned to the hg38 reference genome containing no ALT contigs through bwa-meth (version 0.2.2).^72^ In addition, in order to identify both GpC methylation and CpG methylation, the Biscuit pileup program (version 0.3.16)^73^ was utilized with the parameters “-q 12, -N”. To differentiate exogenous and endogenous DNA methylation, we take advantage of the fact that these modifications occur within different base contexts. Specifically, endogenous DNA methylation occurs in CpG contexts while M.CviPI introduces DNA methylation in GpC contexts. To differentiate the two, we utilize the Biscuit analysis pipeline and specify that only “GCH” are considered for lamina signal detection and only HCG sites are considered for endogenous DNA methylation detection (H=ATC). By only considering GCH and HCG, we exclude GpCpG sites where the methylation signal would be ambiguous. In particular, to obtain position specific information on GpC and CpG methylation, the Biscuit vcf2bed program was employed with the parameters “-k 3” and “-t gch” or “-t hcg” to exclude GpCpG sites, in which methylation signal would be ambiguous. Juicebox (version 1.11.08)^71^ was used for exploratory data analysis and for plotting Hi-C contact maps. The bedgraph files generated from the methylation analysis were converted to bigwig format utilizing the bedGraphToBigWig function (UCSC), which was subsequently imported into Integrated Genome Browser (version 2.5.3)^74^ and Juicebox for visualization or used as input for further analysis. Subsequent data processing was performed in Python 3.7.0 (www.python.org) and R 3.6.1 (www.r-project.org).

### LIMe-Hi-C per-read analysis

Data was processed according to the JuiceMe pipeline and the per-read analysis scripts (https://github.com/aidenlab/JuiceMe/tree/master/Analysis) with modifications detailed below. The Biscuit epiread program with the parameters “-N -m 0 -l 0 -c -u -p” was utilized to create a file detailing both the CpG and GpC methylation status of individual reads from the methylation.bam output of the JuiceMe pipeline (Biscuit, version 0.3.16).^73^ A custom awk script was employed to format the methylation status of the reads appropriately. For the contact matrix file, only intrachromosomal contacts from different fragments with a MAPQ score >0 were considered. After filtering the contact matrix for reads from a chromosome of interest, they were reassigned to a chromosomal value by their methylation status. For example, a read originally mapping to chromosome 1 would be assigned to “chr1m” if it were methylated and “chr1u” if it were unmethylated. A read was considered CpG methylated if 50% of CpG sites within the read were methylated. A read was considered GpC methylated if 10% of GpC sites within the read were methylated. Contacts were then sorted according to their methylation status. To reduce bias from fragments arising from directly neighboring regions, contacts were discarded if the read pair mapped to within less than 1kb of one another. Finally, these contacts were outputted to a hic matrix file using the Juicer tools “pre” function. Matrix files detailing contacts by methylation status were then generated using the Juicer tools “dump” function. The M matrix file contains counts for bins where both read pairs are methylated while the U matrix file contains counts for bins where both read pairs are unmethylated. The Y matrix file is the unsymmetric matrix containing counts for contacts where the read at index i is methylated but the read at index j is not. These files were then loaded into python and analyzed as previously described.^40^

The M matrix for CpG and GpC methylation was directly visualized in Juicebox and displayed in Figure 1. Observed minus expected co-methylation is plotted in heatmap form and is defined as previously described:

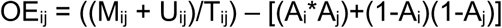

The total number of contacts, T_ij_, is defined as:

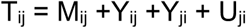

The average methylation of a given locus, A_i_, is defined as:

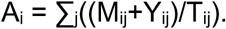

To obtain information on positional dependence of observed methylation across sub-compartments, the observed methylation matrix is defined as:

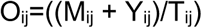

and was min-max normalized by using the average methylation in the A compartment as the min value and the average methylation in the B compartment as the max to correct for differences in dynamic range between samples. The average value of this matrix across 50 kb bins for interactions between compartment or sub-compartments is displayed in heatmap form.

### LIMe-ID Analysis

LIMe-ID analysis was performed analogously to the LIMe-Hi-C analysis except that reads were aligned utilizing the Biscuit alignment pipeline with the following parameters: “biscuit align -t 16 -R ‘@RG\tID’ Read1.fastq.gz Read2fastq.gz | samblaster | samtools sort -o sample.bam -O BAM –” (Biscuit, version 0.3.16).^73^ In addition, to obtain position specific information regarding GpC and CpG methylation, the Biscuit vcf2bed program was employed with the parameters “-k 1” instead of “-k 3” to account for the lower sequencing depth utilized for LIMe-ID.

### Sub-compartment identification and integration of published data

GpC methylation, CpG methylation, and principal component 1 values were determined by averaging across 50 kb bins utilizing the multiBigwigSummary function (deepTools, version 3.4.3). For CpG methylation analysis, CpG islands were excluded by subtracting CpG sites that overlapped with previously annotated islands (UCSC), which were padded by 2 kb using Bedtools2 (version 2.26.0).^75^ These values were *z*-score normalized, and the average values across the three replicates for the DMSO-induced condition were calculated for every 50 kb bin. Regions with an average |*z-*score| > 3 were excluded from the classification. *K*-means clustering and visualization were performed in R (version 3.6.1) with pheatmap (kmeans=4) (version 1.0.12)^76^ across the average *z*-score normalized GpC methylation, CpG methylation, and principal component 1 values to classify each 50 kb genomic bin into a sub-compartment. Published ChIP-seq and DamID data (see Supplementary Table 1) as well as quantitative ChIP-seq data, processed as described below, were similarly averaged across 50 kb bins using the multiBigwigSummary function to determine sub-compartment enrichment and correlation with LIMe-Hi-C features.

### Quantitative ChIP-seq analysis

Reads were aligned to a composite reference genome for dm6 and hg38 utilizing bowtie2 (version 2.3.2)^77^ according to the Spiker analysis framework (version 1.03).^78^ Aligned reads were sorted using samtools (version 0.1.19)^79^ and the Spiker split_bam.py script was run to calculate scaling factors and to split reads into an alignment file containing reads specifically aligning to the hg38 genome (Spiker, version 1.0.3).^78^ This alignment file was then converted into spike-in normalized bigwig file for further analysis and visualization utilizing the deepTools bamCoverage command with the following parameters: -binSize 50 --normalizeUsing RPKM --ignoreDuplicates --ignoreForNormalization chrM --extendReads --scaleFactor. Peaks for individual H3K27me3 replicates were called utilizing MACS2 “callpeak” function with the following parameters: -f BEDPE -B -g hs --broad --broad-cutoff 0.2. Regions within 10kb of one another were merged utilizing the Bedtools2 merge function -d 10000 (Bedtools2, version 2.26.0). Finally, individual replicate peaks were subsequently combined utilizing Bedtools2 merge. For bin level analysis, the H3K27me3 signal value was averaged across 50 kb bins utilizing the multiBigwigSummary function (deepTools, version 3.4.3).^80^ To generate metaprofile plots comparing between treatment conditions the “log2” option in the bigwigCompare function was used prior to plotting with the plotHeatmap function (deepTools, version 3.4.3).^80^

### Identifying LIMe LADs and differential lamina GpC methylated regions

Genome-wide 50 kb GpC methylation data was calculated as specified above and *z*-score transformed. For every replicate, the Bumphunter package (version 1.34.0)^45,46^ in R was subsequently used to identify GpC methylated (*i*.*e*., lamina attached) regions with a threshold of 0. Regions that contained 2 or less contiguous bins (<100 kb) were removed. The set of high confidence LIMe LADs were then defined to be those regions present in at least two of the three replicates. The overlap between LIMe LADs and DamID LADs was calculated for all regions with a sub-compartment designation using pybedtools (version 0.8.1)^81^ “intersect” and “subtract” functions.

To identify differential lamina GpC methylated regions upon either EZH2 or DNMT1 inhibition, a *t*-statistic was calculated by comparing the *z*-score normalized GpC methylation data of three replicates for the inhibitor samples to that of the vehicle-treated samples. A threshold of 4 was utilized to identify differential regions upon EZH2i and threshold of 2 was utilized to determine differential regions upon DNMT1i. Differential regions that contained 2 or less contiguous bins (<100 kb) were excluded from the set of differential regions.

### Principal component analysis and contact frequency analysis

The JuiceMe merged_nodups.txt output file containing a list of paired contacts was reformatted and converted to the HiCSummary format (Homer).^82^ Homer tag directories were generated, and the runHiCpca function with the parameters “-res 50000 -superRes 50000 -std 4 -min 0.25” was used to calculate the first principal component of the Hi-C contact matrix.

To perform the A/B ratio and observed/expected contact frequency analysis, Knight-Ruiz normalized 50 kb binned coordinate sorted matrices were generated using the Juicertools “dump” function. As input compartment designations, the average of principal component 1 across the three replicates was calculated. The bin was assigned to the A compartment if this value was > 0 and assigned to the B compartment if this value was < 0. The observed/expected contact frequency was calculated as previously described.^35^ A similar analysis was performed for the sub-compartment designations except sub-compartment labels were utilized as input regions to calculate average observed/expected contact frequencies.

### H3K27me3-ordered sub-compartment interaction plots

Contact files (.hic) were converted to the cool format using the hic2cool function (version 0.8.3).^83^ Expected contact frequencies were calculated through the following command: “cooltools compute-expected --weight-name KR -t cis” (cooltools, version 0.4.0).^84^ For H3K27me3-ordered sub-compartment interaction plots, 50 kb regions were assigned to a quantile based on H3K27me3 levels within their respective sub-compartment, and these designations were used as input for the following command: “cooltools compute-saddle -t cis --weight-name KR --quantiles -n 100” with chromosomal arms as input regions. These values were log_2_-transformed and averaged over replicates. Standard plotting functions (Python version 3.7.0) were then employed to display the calculated saddle output. To compare inhibitor to vehicle treatment, the difference of the averaged log_2_-transformed saddle data across replicates was calculated and displayed.

### Measuring the distance of regions to borders and their overlap with sub-compartments

Confident B compartment calls were determined through findHiCCompartments (Homer) for all three DMSO plus dox induction replicates with the parameter “-opp”. Regions present in at least two of the three replicates were used to define the set of high confidence B regions. Borders were specified to be the single base pair coordinate at the boundaries of the B compartment domains. Boundary coordinates were specified in a similar manner for the LIMe LADs. The distance of every differential region to a border was determined through the pybedtools (version 0.8.1)^81^ “closest” function, and the median across the whole set of regions was calculated. To determine how this median distance compared to that which would be expected by random chance, the coordinates of the differential regions were shuffled within a chromosome with the pybedtools “shuffle” function, and the same calculation was performed. This random shuffling was repeated 1000 times. An analogous calculation was performed to determine overlap of differential regions with the sub-compartments except that the pybedtools Jaccard metric was utilized in place of the “closest” function. The ratio of the observed Jaccard metric to the simulation average Jaccard metric was calculated to measure the enrichment of a set of regions within a specific sub-compartment.

### Metaprofile plots

aggregate heatmaps were generated utilizing the computeMatrix and plotHeatmap functions (deepTools, version 3.4.3) for specified genomic regions with bigwig files for H3K27me3 and H3K9me3 ChIP-seq enrichment and differential GpC methylation. Differential GpC methylation was calculated by subtracting *z*-score normalized GpC methylation signal averaged across the replicates between vehicle and inhibitor treatments.

The genomic distribution of the sub-compartments was determined by utilizing the computeMatrix and plotProfile functions (deepTools version 3.4.3) with high confidence B compartment and A compartment regions (calculated as specified above for B compartments except using findHiCCompartments without the “-opp” parameter). The presence or absence of a sub-compartment bin at a given location was specified, and regions with no coverage for a sub-compartment were given a zero score for enrichment using the “missingDataAsZero” parameter in the computeMatrix function.

### RNA-seq processing and differential gene expression analysis

RNA-seq data were processed according to the Quantseq 3’ mRNA-Seq Library prep recommended analysis pipeline through alignment to the Ensemble hg38 transcriptome. HTSeq (version 0.11.2)^85^ was used to generate count files. DESeq2 (version 1.28.1)^86^ with R (version 4.0.2) was employed for the differential expression and FPM analysis, and BioMart (version 2.44.1)^87^ with R was used to map genomic ranges of the transcripts. Bedtools2 (version 2.26.0) map was used to annotate genes with sub-compartments, and Bedtools2 coverage was used to determine H3K27me3 status, where a gene was considered H3K27me3-positive if at least 1 base pair overlapped with an H3K27me3 peak. Genes that did not overlap any of the sub-compartments (< 1% of non-zero genes) were excluded from the analysis.

### TRIP expression analysis

Published K562 TRIP data utilized in this study are specified in Supplementary Table 1. Overlap of the promoter integrations with sub-compartments and H3K27me3 peaks was determined through Bedtools2 (version 2.26.0) intersect. The log_2_ transformation of the reported expression value is depicted in the boxplots.

### Statistical methods

Statistical tests and parameters used are reported directly in the figure legends. The definition of center, dispersion and precision measurements (mean +/-s.d. or s.e.m) are reported in the figures and figure legends. The spearman correlation was calculated with the stats.spearmanr function (SciPy, version 1.2.1). The *t*-statistic metric was determined with the two-sided stats.ttest_ind function or the one-sided ttest_1samp function (SciPy, version 1.2.1) assuming unequal variance. For all boxplots, the median and the interquartile range (IQR) are depicted by the box, and the whiskers maximally extend to 1.5 × IQR. Exclusion of outlier points outside this range is specified in the figure legend. Linear regression was performed using the stats.linregress function (SciPy, version 1.2.1) with the R^2^ value specified in the figure. For all plots, replicate-average measurements are employed unless otherwise stated.

## References

1. Lieberman-Aiden, E. et al. Comprehensive Mapping of Long-Range Interactions Reveals Folding Principles of the Human Genome. Science 326, 289–293 (2009).

2. Rao, S., Huntley, M., Durand, N. & Cell, S.-E. A 3D map of the human genome at kilobase resolution reveals principles of chromatin looping. Cell 1665–1680 (2014).

3. Steensel, B. van & Belmont, A. S. Lamina-Associated Domains: Links with Chromosome Architecture, Heterochromatin, and Gene Repression. Cell 169, 780–791 (2017).

4. Kind, J. et al. Genome-wide Maps of Nuclear Lamina Interactions in Single Human Cells. Cell 163, 134–147 (2015).

5. Steensel, B. van & Furlong, E. E. M. The role of transcription in shaping the spatial organization of the genome. Nat Rev Mol Cell Bio 20, 327–337 (2019).

6. Misteli, T. The Self-Organizing Genome: Principles of Genome Architecture and Function. Cell 183, 28–45 (2020).

7. Lochs, S. J. A., Kefalopoulou, S. & Kind, J. Lamina Associated Domains and Gene Regulation in Development and Cancer. Cells 8, 271 (2019).

8. Marchal, C., Sima, J. & Gilbert, D. M. Control of DNA replication timing in the 3D genome. Nat Rev Mol Cell Biology 20, 721–737 (2019).

9. Zheng, H. & Xie, W. The role of 3D genome organization in development and cell differentiation. Nat Rev Mol Cell Bio 20, 535–550 (2019).

10. Kempfer, R. & Pombo, A. Methods for mapping 3D chromosome architecture. Nat Rev Genet 1–20 (2019) doi:10.1038/s41576-019-0195-2.

11. Boettiger, A. N. et al. Super-resolution imaging reveals distinct chromatin folding for different epigenetic states. Nature 529, 418–422 (2016).

12. Bintu, B. et al. Super-resolution chromatin tracing reveals domains and cooperative interactions in single cells. Science 362, eaau1783 (2018).

13. Dekker, J., Rippe, K., Dekker, M. & science, K.-N. Capturing chromosome conformation. Science 295, 1306–1311 (2002).

14. Guelen, L. et al. Domain organization of human chromosomes revealed by mapping of nuclear lamina interactions. Nature 453, 948–951 (2008).

15. Jerković, I. & Cavalli, G. Understanding 3D genome organization by multidisciplinary methods. Nat Rev Mol Cell Bio 22, 511–528 (2021).

16. Rao, S. S. P. et al. Cohesin Loss Eliminates All Loop Domains. Cell 171, 305–320.e24 (2017).

17. Schwarzer, W. et al. Two independent modes of chromatin organization revealed by cohesin removal. Nature 551, 51 (2017).

18. Hildebrand, E. M. & Dekker, J. Mechanisms and Functions of Chromosome Compartmentalization. Trends Biochem Sci 45, 385–396 (2020).

19. Belaghzal, H. et al. Liquid chromatin Hi-C characterizes compartment-dependent chromatin interaction dynamics. Nat Genet 1–12 (2021) doi:10.1038/s41588-021-00784-4.

20. Guerreiro, I. & Kind, J. Spatial chromatin organization and gene regulation at the nuclear lamina. Curr Opin Genet Dev 55, 19–25 (2019).

21. Hansen, K. D. et al. Increased methylation variation in epigenetic domains across cancer types. Nat Genet 43, 768–775 (2011).

22. Berman, B. P. et al. Regions of focal DNA hypermethylation and long-range hypomethylation in colorectal cancer coincide with nuclear lamina–associated domains. Nat Genet 44, 40–46 (2012).

23. Towbin, B. D. et al. Step-Wise Methylation of Histone H3K9 Positions Heterochromatin at the Nuclear Periphery. Cell 150, 934–947 (2012).

24. Hon, G. C. et al. Global DNA hypomethylation coupled to repressive chromatin domain formation and gene silencing in breast cancer. Genome Res 22, 246–258 (2012).

25. Harr, J. C. et al. Directed targeting of chromatin to the nuclear lamina is mediated by chromatin state and A-type lamins. J Cell Biology 208, 33–52 (2015).

26. Kind, J. et al. Single-Cell Dynamics of Genome-Nuclear Lamina Interactions. Cell 153, 178–192 (2013).

27. Meuleman, W. et al. Constitutive nuclear lamina–genome interactions are highly conserved and associated with A/T-rich sequence. Genome Res 23, 270–280 (2013).

28. Zheng, X., Kim, Y. & Zheng, Y. Identification of lamin B–regulated chromatin regions based on chromatin landscapes. Mol Biol Cell 26, 2685–2697 (2015).

29. Zheng, X. et al. Lamins Organize the Global Three-Dimensional Genome from the Nuclear Periphery. Mol Cell 71, 802–815.e7 (2018).

30. Blackledge, N. P., Rose, N. R. & Klose, R. J. Targeting Polycomb systems to regulate gene expression: modifications to a complex story. Nat Rev Mol Cell Bio 16, 643–649 (2015).

31. McLaughlin, K. et al. DNA Methylation Directs Polycomb-Dependent 3D Genome Re-organization in Naive Pluripotency. Cell Reports 29, 1974–1985.e6 (2019).

32. Du, Z. et al. Polycomb Group Proteins Regulate Chromatin Architecture in Mouse Oocytes and Early Embryos. Mol Cell 77, 825–839.e7 (2019).

33. Cai, Y. et al. H3K27me3-rich genomic regions can function as silencers to repress gene expression via chromatin interactions. Nat Commun 12, 719 (2021).

34. Zhang, X. et al. Large DNA Methylation Nadirs Anchor Chromatin Loops Maintaining Hematopoietic Stem Cell Identity. Mol Cell 78, 506–521.e6 (2020).

35. Johnstone, S. E. et al. Large-Scale Topological Changes Restrain Malignant Progression in Colorectal Cancer. Cell 182, 1474–1489 (2020).

36. Kraft, K. et al. Polycomb-mediated Genome Architecture Enables Long-range Spreading of H3K27 methylation. Biorxiv (2020) doi:10.1101/2020.07.27.223438.

37. Kriz, A. J., Colognori, D., Sunwoo, H., Nabet, B. & Lee, J. T. Balancing cohesin eviction and retention prevents aberrant chromosomal interactions, Polycomb-mediated repression, and X-inactivation. Mol Cell (2021) doi:10.1016/j.molcel.2021.02.031.

38. Rhodes, J. D. P. et al. Cohesin Disrupts Polycomb-Dependent Chromosome Interactions in Embryonic Stem Cells. Cell Reports 30, 820–835.e10 (2020).

39. Yu, J.-R., Lee, C.-H., Oksuz, O., Stafford, J. M. & Reinberg, D. PRC2 is high maintenance. Gene Dev 33, 903–935 (2019).

40. Stamenova, E. K. et al. The Hi-Culfite assay reveals relationships between chromatin contacts and DNA methylation state. Biorxiv 481283 (2018) doi:10.1101/481283.

41. Li, G. et al. Joint profiling of DNA methylation and chromatin architecture in single cells. Nat Methods 16, 991–993 (2019).

42. Lee, D.-S. et al. Simultaneous profiling of 3D genome structure and DNA methylation in single human cells. Nat Methods 16, 999–1006 (2019).

43. Xu, M., Kladde, M. P., Simpson, R. T. & Etten, J. L. V. Cloning, characterization and expression of the gene coding for a cytosine-5-DNA methyltransferase recognizing GpC. Nucleic Acids Res 26, 3961–3966 (1998).

44. Carvin, C. D., Dhasarathy, A., Friesenhahn, L. B., Jessen, W. J. & Kladde, M. P. Targeted cytosine methylation for in vivo detection of protein–DNA interactions. Proc National Acad Sci 100, 7743–7748 (2003).

45. Jaffe, A. E. et al. Bump hunting to identify differentially methylated regions in epigenetic epidemiology studies. Int J Epidemiol 41, 200–209 (2012).

46. Aryee, M. J. et al. Minfi: a flexible and comprehensive Bioconductor package for the analysis of Infinium DNA methylation microarrays. Bioinformatics 30, 1363–1369 (2014).

47. Verma, S. K. et al. Identification of Potent, Selective, Cell-Active Inhibitors of the Histone Lysine Methyltransferase EZH2. Acs Med Chem Lett 3, 1091–1096 (2012).

48. Gilmartin, A. G. et al. In vitro and in vivo induction of fetal hemoglobin with a reversible and selective DNMT1 inhibitor. Haematologica haematol.2020.248658 (2020) doi:10.3324/haematol.2020.248658.

49. Pappalardi, M. B. et al. Discovery of a first-in-class reversible DNMT1-selective inhibitor with improved tolerability and efficacy in acute myeloid leukemia. Nat Cancer 2, 1002–1017 (2021).

50. Du, Q. et al. DNA methylation is required to maintain DNA replication timing precision and 3D genome integrity. Biorxiv 2020.10.15.338855 (2020) doi:10.1101/2020.10.15.338855.

51. Spracklin, G. et al. Heterochromatin diversity modulates genome compartmentalization and loop extrusion barriers. doi:10.1101/2021.08.05.455340.

52. Plys, A. J. et al. Phase separation of Polycomb-repressive complex 1 is governed by a charged disordered region of CBX2. Gene Dev 33, 799–813 (2019).

53. Luperchio, T. R. et al. Chromosome Conformation Paints Reveal The Role Of Lamina Association In Genome Organization And Regulation. Biorxiv 122226 (2017) doi:10.1101/122226.

54. Cruz, C. C. de la et al. The Polycomb Group Protein SUZ12 regulates histone H3 lysine 9 methylation and HP1α distribution. Chromosome Res 15, 299–314 (2007).

55. Boros, J., Arnoult, N., Stroobant, V., Collet, J.-F. & Decottignies, A. Polycomb Repressive Complex 2 and H3K27me3 Cooperate with H3K9 Methylation To Maintain Heterochromatin Protein 1α at Chromatin. Mol Cell Biol 34, 3662–3674 (2014).

56. Qi, W. et al. An allosteric PRC2 inhibitor targeting the H3K27me3 binding pocket of EED. Nat Chem Biol 13, 381–388 (2017).

57. Margueron, R. et al. Role of the polycomb protein EED in the propagation of repressive histone marks. Nature 461, 762–767 (2009).

58. Nabet, B. et al. The dTAG system for immediate and target-specific protein degradation. Nat Chem Biol 14, 431–441 (2018).

59. Leemans, C. et al. Promoter-Intrinsic and Local Chromatin Features Determine Gene Repression in LADs. Cell 177, 852–864.e14 (2019).

60. Wutz, G. et al. Topologically associating domains and chromatin loops depend on cohesin and are regulated by CTCF, WAPL, and PDS5 proteins. Embo J 36, 3573–3599 (2017).

61. Haarhuis, J. H. I. et al. The Cohesin Release Factor WAPL Restricts Chromatin Loop Extension. Cell 169, 693–707.e14 (2017).

62. Nuebler, J., Fudenberg, G., Imakaev, M., Abdennur, N. & Mirny, L. A. Chromatin organization by an interplay of loop extrusion and compartmental segregation. Proc National Acad Sci 115, 201717730 (2018).

63. Chen, S. et al. A Lamina-Associated Domain Border Governs Nuclear Lamina Interactions, Transcription, and Recombination of the Tcrb Locus. Cell Reports 25, 1729–1740.e6 (2018).

64. Xiong, K. & Ma, J. Revealing Hi-C subcompartments by imputing inter-chromosomal chromatin interactions. Nat Commun 10, 5069 (2019).

65. Chen, Y. et al. Mapping 3D genome organization relative to nuclear compartments using TSA-Seq as a cytological ruler. J Cell Biology 217, 4025–4048 (2018).

66. Canzio, D., Larson, A. & Narlikar, G. J. Mechanisms of functional promiscuity by HP1 proteins. Trends Cell Biol 24, 377–386 (2014).

67. Onder, T. T. et al. Chromatin-modifying enzymes as modulators of reprogramming. Nature 483, 598–602 (2012).

68. Becker, J. S. et al. Genomic and Proteomic Resolution of Heterochromatin and Its Restriction of Alternate Fate Genes. Mol Cell 68, 1023–1037.e15 (2017).

69. Youmans, D. T., Schmidt, J. C. & Cech, T. R. Live-cell imaging reveals the dynamics of PRC2 and recruitment to chromatin by SUZ12-associated subunits. Gene Dev 32, 794–805 (2018).

70. Liau, B. B. et al. Adaptive Chromatin Remodeling Drives Glioblastoma Stem Cell Plasticity and Drug Tolerance. Cell Stem Cell 20, 233–246.e7 (2017).

71. Durand, N. C. et al. Juicer Provides a One-Click System for Analyzing Loop-Resolution Hi-C Experiments. Cell Syst 3, 95–98 (2016).

72. Pedersen, B. S., Eyring, K., De, S., Yang, I. V. & Schwartz, D. A. Fast and accurate alignment of long bisulfite-seq reads. Arxiv (2014).

73. Shen, H. BISulfite-seq CUI Toolkit (BISCUIT). https://huishenlab.github.io/biscuit/.

74. Thorvaldsdóttir, H., Robinson, J. T. & Mesirov, J. P. Integrative Genomics Viewer (IGV): high-performance genomics data visualization and exploration. Brief Bioinform 14, 178–192 (2013).

75. Quinlan, A. R. & Hall, I. M. BEDTools: a flexible suite of utilities for comparing genomic features. Bioinformatics 26, 841–842 (2010).

76. Kolde, R. pheatmap: Pretty heatmaps. https://cran.r-project.org/web/packages/pheatmap/index.html.

77. Langmead, B. & Salzberg, S. L. Fast gapped-read alignment with Bowtie 2. Nat Methods 9, 357–359 (2012).

78. Wu, D., Wang, L. & Huang, H. Protocol to apply spike-in ChIP-seq to capture massive histone acetylation in human cells. Star Protoc 2, 100681 (2021).

79. Li, H. et al. The Sequence Alignment/Map format and SAMtools. Bioinformatics 25, 2078–2079 (2009).

80. Ramírez, F. et al. deepTools2: a next generation web server for deep-sequencing data analysis. Nucleic Acids Res 44, W160–W165 (2016).

81. Dale, R. K., Pedersen, B. S. & Quinlan, A. R. Pybedtools: a flexible Python library for manipulating genomic datasets and annotations. Bioinformatics 27, 3423–3424 (2011).

82. Heinz, S. et al. Transcription Elongation Can Affect Genome 3D Structure. Cell 174, 1522–1536.e22 (2018).

83. Vitzthum, C., Abdennur, N., Lee, S. & Kerpedjiev, P. hic2cool. https://github.com/4dn-dcic/hic2cool.

84. Abdennur, N. & Mirny, L. A. Cooler: scalable storage for Hi-C data and other genomically labeled arrays. Bioinformatics (2019) doi:10.1093/bioinformatics/btz540.

85. Anders, S., Pyl, P. T. & Huber, W. HTSeq—a Python framework to work with high-throughput sequencing data. Bioinformatics 31, 166–169 (2015).

86. Love, M. I., Huber, W. & Anders, S. Moderated estimation of fold change and dispersion for RNA-seq data with DESeq2. Genome Biol 15, 550 (2014).

87. Durinck, S., Spellman, P. T., Birney, E. & Huber, W. Mapping identifiers for the integration of genomic datasets with the R/Bioconductor package biomaRt. Nat Protoc 4, 1184–1191 (2009).

